# CXCL9-Dependent Immune Networks of Spontaneous Tertiary Lymphoid Structures in a Murine Subcutaneous Tumor Model

**DOI:** 10.64898/2026.10.12.758663

**Authors:** Raymond Z. Qin, Hanna Abberger, Guigui Man, Benjamin J. Broomfield, Bailey E. Williams, Chin Wee Tan, Brigette C. Duckworth, Misty R. Jenkins, Niall D. Geoghegan, Kelly L. Rogers, Zewen Kelvin Tuong, Joanna R. Groom

**Affiliations:** Walter and Eliza Hall Institute of Medical Research, Parkville, Victoria, Australia; Department of Medical Biology, University of Melbourne, Parkville, Victoria, Australia; Ian Frazer Centre for Children’s Immunotherapy Research, Child Health Research Centre, Faculty of Health, Medicine and Behavioural Sciences, The University of Queensland, South Brisbane, Queensland, Australia; Drug Delivery, Disposition and Dynamics, Monash Institute of Pharmaceutical Sciences, Monash University, Parkville, Victoria, Australia

## Abstract

Recent discoveries have identified human tumor tertiary lymphoid structures (TLS) as organized clusters of innate and adaptive immune cells and stromal cells. Cancer-associated TLS can promote tumor clearance and enhance responses to immune checkpoint blockade (ICB) immunotherapy. However, the absence of TLS in conventional mouse tumor models, particularly subcutaneous transplant models, limits experimental manipulation and study of TLS development. By tuning tumor antigen expression, we have generated a subcutaneous tumor model that naturally forms TLS within the immune-hot, vasculature-rich peritumoral adipose tissue. These TLS were identified as clusters of B and T cells in close proximity to high endothelial venules. TLS organization was underpinned by spatially compartmentalized chemokine expression, with CCL21 produced in the T cell zone of perivascular TLS and CXCL13 present in the B cell region. Single cell transcriptomic sequencing showed that the immune composition in the TLS forming peritumoral tissue resembled human cancer TLS signatures. Upon anti-PD-L1 ICB treatment, TLS underwent expansion, which was associated with increased interferon responses in macrophages and coincided with tumor clearance. We further identified the interferon-inducible chemokine CXCL9 as a key regulator of the TLS microenvironment. CXCL9-expressing macrophages populated TLS, interacting with B cells following ICB treatment. CXCL9 deficiency reduced B cell recruitment to peritumoral tissue and TLS formation. Further T and B cell receptor sequencing indicated a CXCL9-dependent promotion of antigen-specific responses within the tumor and TLS niche. Combined, these findings establish a tractable subcutaneous model of naturally arising tumor TLS and define the cellular and molecular networks that regulate their formation and function. This provides a platform to dissect how TLS develop and test strategies that promote TLS formation to enhance cancer therapy.

## INTRODUCTION

Tertiary lymphoid structures (TLS) are organized immune aggregates that develop in non-lymphoid tissues at sites of chronic inflammation.^1,2^ TLS have been recognized in human cancers for more than a decade and comprise of immune and stromal cells producing cytokines and chemokines that coordinate their development and function.^3,4^ Although they arise outside of conventional lymphoid organs, TLS recapitulate key features of secondary lymphoid organs such as the lymph nodes and can generate and sustain local immune responses against tumor antigens.^5^ Importantly, the presence of TLS has emerged as a favorable prognostic marker for some solid tumors and is often associated with improved outcomes following immunotherapy.^6–8^ Defining the cellular and molecular networks that govern TLS formation and function, and understanding how they promote therapeutic responses, is an essential step towards harnessing them to enhance anti-tumor immunity.

Human TLS are classified both by their anatomical regions in the tumor and by maturation stages.^9^ Immature TLS contain segregated B and T cell zones and developing lymphatic and stromal networks, while mature TLS contain well-developed germinal centers (GC) within the B cell follicles, to support B cell affinity maturation and the generation of antibody-secreting cells.^5,6,10^ Within TLS, B cells can also act as antigen presenting cells to promote T cell activation and improve cancer outcomes.^7,11,12^ Notably, TCF-1^+^ CD8^+^ stem-like T (T_SL_) cells, a self-renewable memory T cell subset capable of generating CD8^+^ effector T cells, have been observed within TLS.^7,13–16^ However, other studies report that T_SL_ cells reside in distinct immune hubs rather than bonafide TLS.^17–20^ This discrepancy raises the question of whether TLS are required for the local maintenance of T_SL_ cells within the tumor microenvironment. Among CD4^+^ T cells, follicular helper T (T_FH_) cells are present in TLS and found to interact with follicular and GC B cells.^21^ Together, these observations suggest that the recruitment and interactions of these immune populations drive TLS neogenesis and function. Immunotherapy, such as immune checkpoint blockade (ICB), can remodel TLS^22^ and promote the infiltration of B cells,^23^ T_SL_,^24^ and T_FH_,^25^ thereby amplifying anti-tumor immunity. However, most studies in humans have been correlative and retrospective, leaving the direct mechanisms poorly understood. A central challenge is understanding how immune cell states and TLS biology jointly shape responses to ICB.

Chemokine expression is a critical feature within the tumor microenvironment.^26,27^ Multiple chemokine axes, including CCR7, CXCR3, CXCR4, and CXCR5, direct immune cell recruitment and shape cellular interactions in cancer. Lymphoid chemokines CCL19/21, CXCL12, and CXCL13 coordinate the recruitment and spatial organization of dendritic cells (DC), T cells and B cells within TLS.^28–32^ In melanoma, CXCL9-expressing cells localize to immune hubs enriched in CXCL13 expression,^19^ suggesting the potential co-recruitment of B cells and T cells. In mouse cancer models, CXCL9 expression by lymphatic vessels can promote immune cell recruitment,^33,34^ and can facilitate interactions between CXCR3 expressing CD8^+^ T cells and CXCL9^+^ type 1 conventional DCs (cDC1), thereby supporting effective immunotherapy responses.^35,36^ Conversely, CXCR3^+^ CD4^+^ regulatory T (T_REG_) cells impede anti-tumor responses.^37^ Together, these findings suggest that CXCL9 functions in a context-dependent manner within the tumor microenvironment, with its effects influenced by the local niche and the immune cells it recruits.

Across many human solid tumor types, TLS can develop within the tumors, in the adjacent peritumoral tissue or in the distal tissue.^9^ Particularly, the peritumoral adipose stromal tissue is proposed to form an immune-hot, vascularized niche that facilitates immune cell recruitment.^31,38–42^ However, only a limited number of mouse models recapitulate this TLS development.^32,43–46^ Many commonly used models, including transplanted cell lines and spontaneous models exhibit limited immune infiltration and represent the “immune desert” archetype of human cancers.^47^ Subcutaneous transplantation of tumor cell lines is widely used because this tumor growth can be readily monitored. However, these tumors grow rapidly and resemble an acute, overwhelming response, rather than establishing a sustained tumor-immune equilibrium. We therefore reasoned that tuning tumor antigen expression could slow tumor growth, permitting sustained immune cell interactions and TLS formation within the tumor microenvironment.

Here, we describe a subcutaneous tumor model with intermediate immunogenicity that supports the formation of organized, immature TLS. Since human colorectal cancer is among the highest TLS-forming solid cancers,^48^ we engineered a Murine Colon Adenocarcinoma 38 (MC38) cell line-based transplantation model, providing an interpretive bridge from a widely used mouse model to a clinically TLS-relevant setting. To our knowledge, this represents the first description of a subcutaneous tumor model with spontaneous TLS formation. The identification of TLS within the peritumoral adipose tissue niche enabled mechanistic dissection of TLS biology and their contribution to anti-tumor immunity. This model recapitulates key cellular features of human cancer TLS, including memory CD8^+^ and CD4^+^ T cells and naïve-like B cells, and is characterized by spatially organized chemokine expression. Building on the reported association between TLS and improved ICB responses in human cancers, we show that the TLS niche was the primary site of the response to anti-PD-L1 ICB and supported immune cell recruitment and interactions that promoted effective tumor control. ICB elicited an interferon-CXCL9 signature in TLS, and CXCL9 was required for immunotherapy-driven antigen-specific responses within the tumor and TLS niche. Together, development and characterization of this model provides a mechanistic understanding of TLS function and provides avenues to therapeutically promote TLS formation to enhance cancer therapy.

## RESULTS

### Establishing a murine model of spontaneous subcutaneous peritumoral TLS

Human colorectal cancer is one of the highest TLS-forming solid cancers.^48^ To investigate how titrating neoantigen expression impacts tumor growth, we used the MC38 colorectal carcinoma cell lines engineered with distinct levels of the model antigen ovalbumin (OVA) expression. This included the wildtype (WT) MC38 cell line without OVA expression (MC38-WT), MC38 cells transduced to express intermediate/low OVA expression (MC38-OVA^lo^), and MC38 cells transduced to express high OVA expression (MC38-OVA^hi^). The level of the OVA immunodominant epitope SIINFEKL presented in the context of MHC-I (H-2K^b^-SIINFEKL) showed that both MC38-OVA^lo^ and MC38-OVA^hi^ cell lines had high frequencies of OVA expression, with the latter having higher mean fluorescent intensity of antigen presentation (Figure. 1A). Each of these three MC38 cell lines were subcutaneously injected into the left flank of WT C57BL/6 mice and tumor growth was measured longitudinally. The higher immunogenic MC38-OVA^hi^ tumor model reduced the growth rate and achieved tumor clearance in some mice, while the MC38-OVA^lo^ model displayed gradual growth compared to MC38-WT (Figure. 1B). To assess how differences in tumor growth impacted the tumor immune microenvironments, tumors were analyzed by confocal imaging and flow cytometry on day 15 post tumor transplantation. MC38-WT and MC38-OVA^lo^ cell lines exhibited comparable immune cell infiltration and density in tumors, while this was increased in the MC38-OVA^hi^ model (Figure. 1C,D). Immune cell analyses showed the overall proportion of T and B cells among the immune infiltrates were similar between models, while MC38-OVA^hi^ tumors exhibited the highest proportion of T_REG_ and exhausted CD8^+^ T cells (T_EX_) and lowest proportion of T_FH_-like cells, suggesting a discrepancy between increased immune infiltration and functionality (Figures. S1A-D). Importantly, the MC38-OVA^lo^ tumor model presented with a gradual, sustained tumor growth and functional immune engagement.

**Figure 1.**
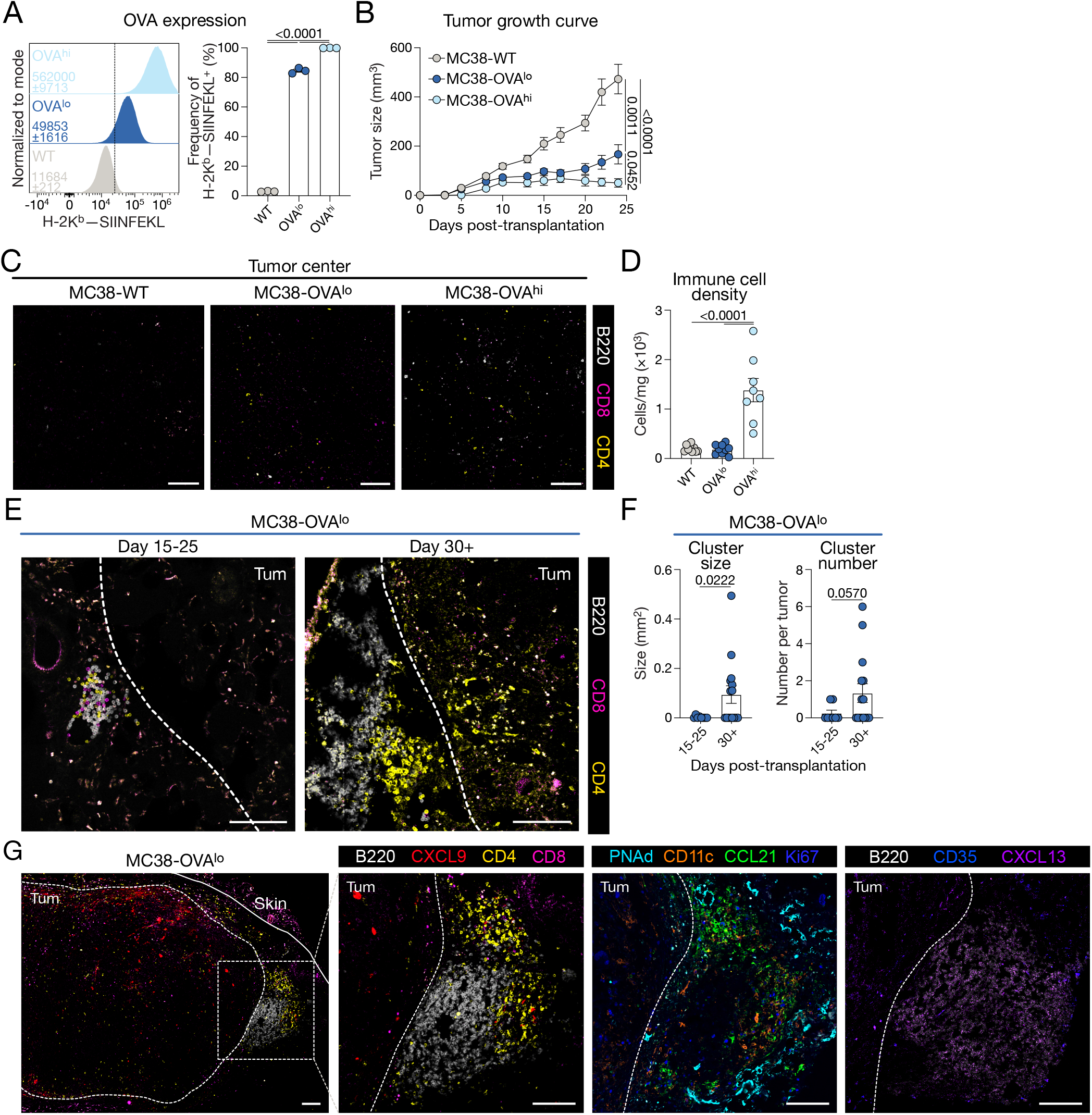
MC38-OVA^lo^ subcutaneous cancer model permits peritumoral TLS formation. (A) Representative histograms of H-2K^b^-SIINFEKL antibody staining for MC38-WT, MC38-OVA^lo^ and MC38-OVA^hi^ cell lines with average geometric mean fluorescent intensity (gMFI) ± SEM indicated. Frequency of OVA-expressing cells for the three MC38 cell lines. *n* =3 per group. (B) Tumor growth curves for subcutaneously transplanted MC38-WT, MC38-OVA^lo^ and MC38-OVA^hi^ tumors. *n* = 13 for MC38-WT, *n* = 13 for MC38-OVA^lo^, *n* = 8 for MC38-OVA^hi^. (C) Representative confocal images of tumor center of indicated tumors on day 15 post transplantation. Scale bars represent 100 µm. (D) Immune cell densities (total number of immune cells/total mass of tumor) from tumors on day 15 post transplantation analyzed by flow cytometry. *n* = 8-9 per group. (E-G) MC38-OVA^lo^ cells were subcutaneously injected and tumors were harvested on days (E-F) 15-25 or 30+ as indicated or (G) day 30 post transplantation. (E) Representative confocal images of MC38-OVA^lo^ peritumoral immune clusters at day 15-25 and day 30+ timepoints. White dashed lines represent tumor border. Tum indicates tumor region. Scale bars represent 100 µm. (F) Quantifications of immune cluster size and number in (E). *n* = 8 for day 15-25, *n* = 15 for day 30+ timepoints. (G) Representative confocal images of serial sections of MC38-OVA^lo^ peritumoral TLS. White dashed lines represent tumor borders. Solid white line represents skin. White dashed square in the left image represents the field the view of subsequent images on the right. Tum indicates tumor region. Scale bars represent 100 µm. Data are mean ± SEM. Statistics for (A), (B), (D) was determined using unpaired one-way ANOVA test. Statistics for (F) was determined using unpaired *t*-tests.

To more thoroughly characterize the tumor immune microenvironment of the MC38-OVA^lo^ model, we conducted serial section confocal imaging. To preserve the overall tissue integrity, tumors were harvested with adjacent peritumoral tissues, including the skin, adipose tissue and peritoneal membrane. We compared the immune infiltration in two experimental windows day 15 to 25 (when an overt tumor mass is evident) and day 30 or more (when tumor growth stabilizes) following transplantation. At day 15-25, we observed the spontaneous formation of lymphoid aggregates, containing B cells, and CD4^+^ and CD8^+^ T cells, exclusively in the peritumoral region of the tumor (Figure. 1E). By day 30+, around 50% of tumors naturally formed more organized immune clusters with B and T cells within the peritumoral region, supporting the notion that this niche provides permissive conditions for immune engagement (Figures. 1E,F).^38,40,41,43^ As human TLS are underpinned by chemokine expression, we next characterized inflammatory and lymphoid chemokine expression in the context of the distinct T and B cell zones, DCs and lymphatic vasculature to determine if these organized immune clusters were bona fide TLS (Figure. 1G). CXCL9 was expressed in the tumor core and T cell zone, CCL21 was associated with the T cell zone while CXCL13 was present in the B cell zone. CD11c^+^ DCs were preferentially located within T cell zones, while the entire structure was surrounded by PNAd^+^ high endothelial venues. Although some proliferative Ki67^+^ and CD35^+^ follicular dendritic cells were observed sparsely throughout and surrounding the structure, no mature GC structure was observed. Therefore, transplantation of MC38-OVA^lo^ tumors led to the establishment of a cancer model that spontaneously forms TLS that structurally resemble human immature TLS.^48,49^

### TLS represent an intermediate immune niche between the draining lymph node and tumor core

As TLS in MC38-OVA^lo^ tumors were identified in the peritumoral tissue, this enabled macroscopic separation of the peritumoral adipose tissue (PAT) from the tumor core (Tum) (Figure. 2A; Figure S2A). This allowed analysis of the PAT immune composition, compared to both the tumor core and tumor draining lymph nodes (dLN). To investigate TLS composition longitudinally, we performed time course experiments, using a spectral flow cytometry panel to assess PAT, Tum and dLN tissues on day 20, 30 and 40 post tumor transplantation. While the total immune cell numbers in the PAT and Tum increased over time, the cell number ratio between tissues peaked at day 30 (Figure. S2B). Broad proportions of B and T cell populations tended to decrease, suggesting an increase in myeloid cell infiltration as tumors progressed (Figures. S2C-F).

**Figure 2.**
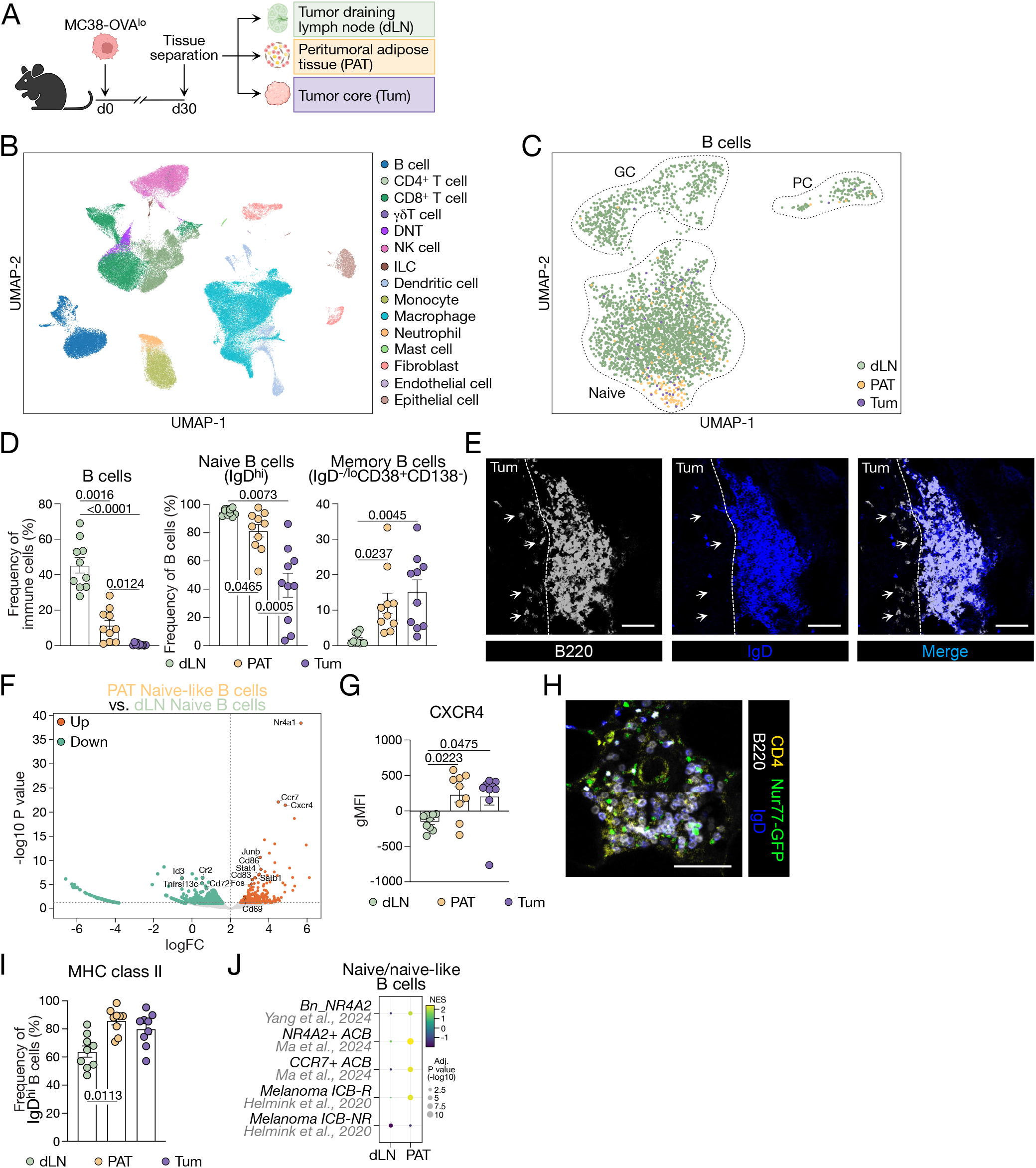
TLS harbor naïve-like B cells. MC38-OVA^lo^ cells were subcutaneously injected and tumors were harvested on day 30 post transplantation. (A) Experimental scheme. (B) UMAP of 134,519 cells from scRNA-seq analysis, colored by 15 broad cell types. DNT: double negative T cell; NK: natural killer; ILC: innate lymphoid cell. (C) UMAP of B cells reclustered from (B) (*n* = 4 mice; 3,347 cells), colored by tissue of origin. Dashed outlines indicate B cell populations. Naïve: naïve B cells; GC: germinal center B cells; PC: plasma cells. (D) Frequencies of B cell subsets analyzed by flow cytometry. *n =* 10 per group. (E) Representative confocal images of MC38-OVA^lo^ peritumoral TLS. White dashed lines represent tumor borders. Tum indicates tumor region. White arrows indicate intratumoral B220^+^ IgD^-^ B cells. Scale bars represent 50 µm. (F) Pseudobulk differential expression analysis comparing PAT naïve B cells with lymph-node naïve B cells, performed using the edgeR quasi-likelihood framework. Positive logFC values indicate higher expression in PAT B cells. Differentially expressed genes were defined at false discovery rate (FDR) < 0.05, yielding 152 significant genes, including 89 upregulated and 63 downregulated genes in PAT B cells. Colored points highlight selected genes of interest, whereas all other genes are shown in grey. Dotted lines indicate *P* = 0.05 and logFC = 0. (G) gMFI of CXCR4 for IgD^hi^ B cells in three tissues. *n* = 9 per group. (H) Representative confocal images of TLS in Nur77-GFP host. Scale bar represents 50 µm. *n* = 3. (I) Frequency of MHCII^+^ for IgD^hi^ B cells in three tissues. *n* = 9 per group. (J) Normalized enrichment score (NES) plot of enrichment of tumor infiltrating B cells from published human datasets. *Bn*: B naïve; *ACB*: activated B cell; *ICB-R/-NR*: immune checkpoint blockade responsive/non-responsive patients. Data are mean ± SEM. Statistics for (D), (F), (G), (I) was determined using paired one-way ANOVA test.

To further dissect the cellular and molecular components of these microenvironments, we performed single cell RNA-sequencing (scRNA-seq) on cells within the Tum and PAT and cells enriched for activated lymphocytes in dLN (Figure. 2A; Figures. S3A,B). A total of 15 clusters and 134519 cells were identified (Figure. 2B; Figure. S3C). DC and macrophage population analysis highlighted an increase in cDC2s in the PAT and tumor core relative to the dLN (Figure. S3D). While macrophage populations were absent in the dLN, a higher proportion of migratory macrophages were found in PAT (Figure. S3E). We next focused on B cell analysis and reclustered the B cells in this dataset. Consistent with the absence of mature TLS, PAT B cells clustered with naïve B cells from the dLN (Figure. 2C). This was confirmed by flow cytometry to demonstrate that peritumoral B cells were predominantly IgD^+^ naïve-like B cell, while intratumoral B cells had a higher percentage of activated IgD^low/-^ B cell (Figure. 2D). Spatially, confocal imaging showed B cells within the peritumoral TLS were mostly IgD^+^, while those within tumor border showed no IgD staining (Figure. 2E). Within the activated B cell population, there was a higher proportion of memory B cells in the PAT and Tum, compared to the dLN (Figure. 2D). To determine how PAT IgD^+^ B cells differ from the truly naïve lymph node B cells, differential gene analysis between PAT naïve B cells and naïve B cells from the dLN was performed. Here, PAT B cells had higher expression of chemokine receptors (*Ccr7*, *Cxcr4*), activation (*Nr4a1, Junb*, *Fos*, *Stat4*) and costimulatory markers (*Cd83*, *Cd86*) (Figure. 2F). Gene expression of CXCR4 was validated at protein level by flow cytometry (Figure. 2G). The transcription factor *Nr4a1* (encoding Nur77) that indicates early B cell receptor (BCR) signaling was elevated in PAT B cells (Figure. 2F). We thus transplanted MC38-OVA^lo^ tumors in Nur77-GFP reporter host mice and identified *Nr4a1^+^* B cells within the PAT TLS (Figure. 2H). We also observed an increased percentage of major histocompatibility complex class II (MHCII)^+^ B cells in the PAT (Figure. 2I), indicating enhanced capacity of antigen presentation and costimulation in the peritumoral TLS niche compared to dLN. Thus, we termed these B cells - naïve-like B cells. Next, to determine whether these naïve-like B cells within the PAT correlated with human TLS or tumor-infiltrating B cells, we compared our gene signature to human published datasets (Figure. 2J). PAT naïve-like B cell gene signatures showed higher similarities with tumor infiltrating B cells in humans and correlated with the gene signatures expressed by ICB responsive TLS positive patient biopsies.^6^ Combined, TLS B cells found in the peritumoral niche of MC38-OVA^lo^ tumors resemble cellular and molecular markers consistent to human cancers.

T_SL_ cells and T_FH_ cells are two key T cell subsets that have been correlated with TLS composition in humans.^15,18,50^ To investigate the T cell composition in MC38-OVA^lo^ TLS, we first clustered CD4^+^ T cells in our scRNA-seq dataset (Figure. 3A). Differential gene expression of PAT and Tum CD4^+^ T cells showed CD4^+^ T cells within PAT expressed higher memory and stem-like cell associated genes (*Tcf7*, *Lef1*, *Foxp1*, *Klf2*, *Bach2*) and lower exhaustion and regulatory genes (*Ctla4*, *Tigit*, *Lag3*, *Foxp3*) (Figure. 3B). Flow cytometry analysis further showed a lower T_REG_ cell proportion in the PAT, while there was no significant change in Ly6C^+^CD4^+^ effector T cells (Figure. 3C). Since the collagenase IV digestion used to dissociate tumor and PAT tissue cleaves CXCR5,^13^ we used PD-1 to identify T_FH_-like cells, however this was variable between tissue and may also indicate cell exhaustion (Figure. 3C). We next clustered CD8^+^ T cells to assess differential gene expression relative to tumor-associated tissue (Figures. 3D,E). Similar to CD4^+^ T cells, CD8^+^ T cells within the PAT showed higher memory and stem-like T cell markers (*Ccr7, Sell, Lef1, Satb1, Il7r, Tcf7*) compared to Tum CD8^+^ T cells, which expressed higher levels of exhaustion markers (*Lag3, Cxcr6, Tigit, Tox, Havcr2*) (Figure. 3E). Using flow cytometry, we identified a higher frequency of CXCR6^+^ T_EX_ cells in the tumor core compared to the PAT and dLN (Figure. 3F). In contrast, TCF-1^+^PD-1^-^ central memory CD8^+^ T cells (T_CM_) were increased in the PAT, while TCF-1^+^PD-1^+^ T_SL_ cells were similar in each niche (Figure. 3F). T cell receptor (TCR) clonal sequencing with inferred pseudotime analysis suggested a differentiation transition from naïve to memory to exhausted along a dLN, PAT, tumor core trajectory as individual clones aligned with distinct cell states from each tissue (Figures. 3G,H). Next, to determine whether these T cell gene signatures could be identified in human cancers, we compared the PAT CD4^+^ and CD8^+^ T cell signatures to published human pseudobulk datasets (Figure. 3I). The PAT T cell signatures correlated with both high presence of TLS and positive response to ICB in human hepatocellular carcinoma. Combined, this analysis demonstrates that the PAT niche, where TLS are present, represents a transitional environment between the dLN and tumor core. Further, B and T cells within the TLS niche, in the MC38-OVA^lo^ model, aligned with human TLS gene signatures.

**Figure 3.**
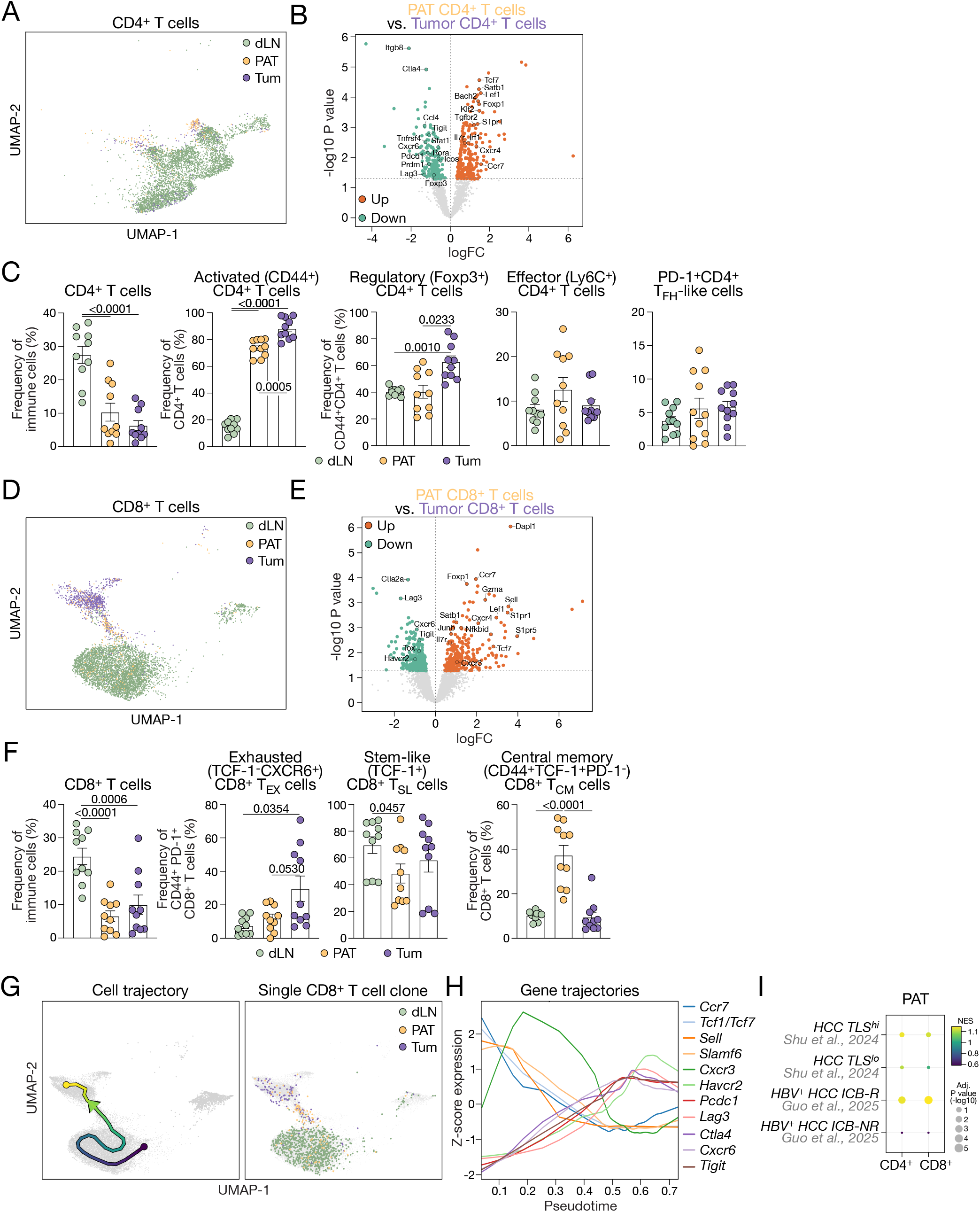
TLS niche exhibits higher memory and lower inhibitory T cells. MC38-OVA^lo^ cells were subcutaneously injected and tumors were harvested on day 30 post transplantation. (A) UMAP of CD4^+^ T cells from Figure. 2A, colored by tissue of origin. (B) Pseudobulk differential expression analysis comparing PAT CD4^+^ T cells with Tum CD4^+^ T cells, performed using the edgeR quasi-likelihood framework. Positive logFC values indicate higher expression in PAT CD4^+^ T cells. Differentially expressed genes were defined at FDR < 0.05, yielding 27 significant genes, including 19 upregulated and 8 downregulated genes in PAT CD4^+^ T cells. Colored points highlight selected genes of interest, whereas all other genes are shown in grey. Dotted lines indicate nominal *P* = 0.05 and logFC = 0. (C) Frequencies of CD4^+^ T cell subsets analyzed by flow cytometry. *n =* 10 per group. (D) UMAP of CD8^+^ T cells, colored by tissue of origin. (E) Paired pseudobulk differential expression analysis comparing PAT CD8^+^ T cells with Tum CD8^+^ T cells. Eight pseudobulk libraries were analysed using the edgeR quasi-likelihood framework. After filtering, 7,513 genes were retained for testing, of which 17 were significantly differentially expressed at FDR < 0.05, including 13 upregulated and 4 downregulated genes in PAT CD8^+^ T cells. Colored points highlight selected genes of interest, whereas all other genes are shown in grey. (F) Frequencies of CD8^+^ T cell subsets analyzed by flow cytometry. *n =* 10 per group. (G) Differentiation trajectory analysis of CD8^+^ T cells shown on the CD8^+^ T cell UMAP. Left, an elastic principal curve fitted using scFates and colored by pseudotime, with arrows indicating the inferred direction of progression. Right, distribution of 1,127 cells belonging to the representative TCR clonotype cluster, colored by tissue of origin. (H) Key CD8^+^ T cell gene expressions along pseudotime. (I) NES plot of enrichment of ICB responsive (R)/non-responsive (NR) and TLS high (hi)/low (lo) gene signatures from human hepatocellular carcinoma cancers. *HCC*: hepatocellular carcinoma; *HBV*: hepatitis B virus. Data are mean ± SEM. Statistics for (C), (F) was determined using paired one-way ANOVA test.

### ICB activity occurs in the TLS

Across multiple human cancers, TLS presence is associated with improved responses to ICB, and ICB may act to enhance existing TLS formation and maturation.^6,9,51^ We therefore tested the effect of anti-PD-L1 immunotherapy on existing immature TLS in the MC38-OVA^lo^ tumor model. Following administration of three doses of anti-PD-L1 from day 22, tumor core and PAT were harvested on day 30 for analysis (Figure. 4A). As anticipated, MC38-OVA^lo^ tumors were responsive to ICB therapy, with tumor growth being controlled relative to control antibody (Figure. 4B). TLS found in the peritumoral niche were remodeled, with increased in size and number with ICB and exhibited a higher degree of T cell infiltration within the B cell zone (Figures. 4C,D). Flow cytometry analyses further demonstrated a significant increase in B cell and CD4^+^ T cell proportion in the PAT, and trend of increase of CD8^+^ T cells (Figure. 4E). Importantly, this activity was confined to the PAT region, as B cell and CD4^+^ T cell percentages remained similar in the tumor core regardless of treatment (Figure. 4E). Within populations of B cells and T cells, differences were observed between tissue sites, such as a decrease in T_EX_ and T_REG_ cells as well as increased T_CM_ cell and naïve-like B cell proportions in the PAT compared to the tumor core (Figures. S4A-C). However, cell proportions were conserved between both ICB treated and non-treated samples (Figures. S4A-C). Together, this showed that anti-PD-L1 ICB acts to increase overall B cell and T cell accumulation in the PAT, correlating with increased TLS formation and better tumor protection, rather than promoting dynamic shifts in immune cell states. To investigate the changes in these microenvironments following ICB treatment, we asked which signaling pathways underpin cellular communication in PAT and Tum B cells. Here, we performed a receptor-ligand analysis on TLS related chemokines and cytokines that may regulate cell recruitment and survival. We observed a reduction of *Dpp4*-chemokine interactions in the PAT, especially in the anti-PD-L1 treated PAT (Figure. 4F). Since DPP4 cleaves multiple chemokines,^52–55^ the loss of this interaction may indicate increased presence of chemokines within the microenvironment, as suggested by the increased immune cell infiltration and increased lymphocyte and antigen presenting cell interactions (Figure. 4E; Figure. S4D). These results suggested that ICB treatment supports anti-tumor immunity by immune cell recruitment into PAT and may facilitate increased interactions within the TLS niche and tumor microenvironment.

**Figure 4.**
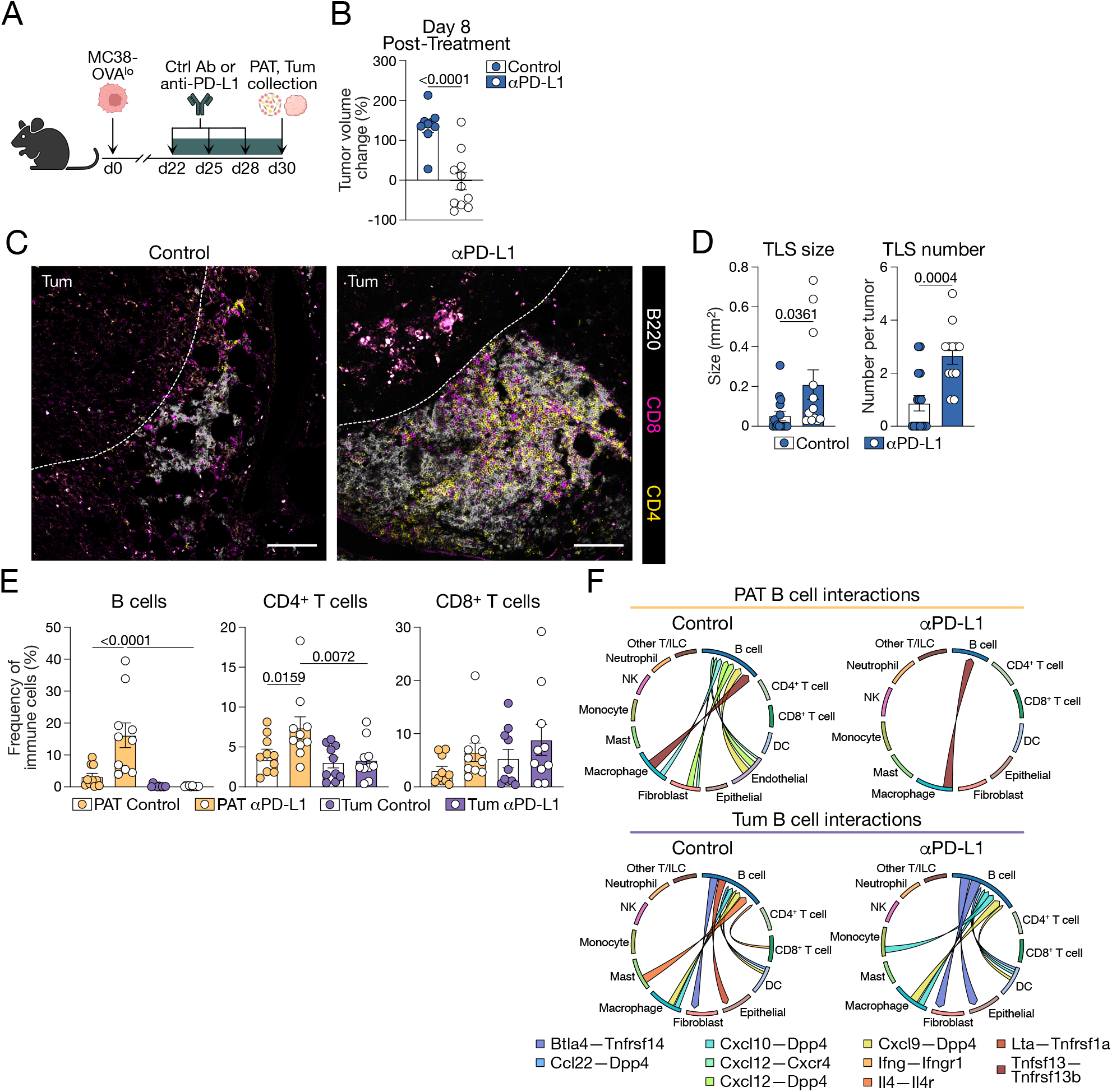
Activity of immune checkpoint blockade occurs in TLS. MC38-OVA^lo^ tumors were subcutaneously injected and, from day 22 post tumor transplantation, treated with three doses of intraperitoneal injected anti-PD-L1 (labeled αPD-L1 on figures) prior to day 30 harvest. (A) Experimental scheme. (B) Relative tumor size change 8 days post anti-PD-L1 treatment. *n* = 9-11 per group. (C) Representative confocal images of control or anti-PD-L1 treated MC38-OVA^lo^ peritumoral TLS. White dashed lines represent tumor border. Tum indicates tumor region. Scale bars represent 100 µm. (D) Quantifications of TLS size and number in (C). *n* = 9-11 per group. (E) Frequencies of B cells, CD4^+^ T cells, CD8^+^ T cell in PAT and Tum with control or anti-PD-L1 treatment. *n* = 10 per group. (F) B cell-associated cytokine/chemokine ligand–receptor interactions in PAT and Tum. Chord diagrams show significant ligand–receptor (L–R) interactions involving B cells detected by CellPhoneDB v5 using the statistical method across four conditions. Sectors represent 13 broad cell-type populations. Ribbons are colored by L–R pair according to the shared legend across all four panels, and interaction direction is shown from the ligand-producing cell type to the receptor-expressing cell type. NK: natural killer; ILC: innate lymphoid cell. Data are mean ± SEM. Statistics for (B) and (D) was determined using unpaired *t*-test. Statistics for (E) was determined using two-way ANOVA test.

### ICB induces a transcriptional switch in macrophages within TLS

The observation of ICB-induced dynamic cellular and molecular changes prompted us to perform gene set enrichment analysis of PAT with or without ICB treatment. A dominant ICB response was the upregulation of interferon-α/β (IFNα, IFNβ) and interferon-γ (IFNγ) response pathways observed in macrophages, monocytes and S100a4^+^ fibroblasts (Figure. 5A). Since the stimulation of type I and II IFN pathways can induce the expression of TLS-forming factors and are enriched in TLS forming regions,^56–59^ we next investigated the potential mechanisms of increased TLS formation in our model. Differential gene expression within macrophages demonstrated higher antigen presentation (*H2-M3, H2-D1, Ciita*) and activation genes (*Irf8, Stat1, Irf1, Fcgr4*) as well as expression of the IFN inducible chemokine CXCL9 (Figure. 5B). We identified a switch in macrophage phenotype from high *Spp1* expression to high *Cxcl9* expression upon ICB treatment (Figure. 5C). This correlates with a recent human study that identified the CXCL9:SPP1 axis as a key macrophage predictor of tumor outcomes.^60^ The macrophage signature generated in our mouse model was aligned with this human dataset (Figure. 5D). To understand how CXCL9-producing cells are spatially distributed in the TLS and tumor core, we used CXCL9 chemokine reporter REX3 mice.^61^ Following anti-PD-L1 treatment, CXCL9-producing F4/80^+^ macrophages were found occupying the TLS niche in close proximity to B cells (Figure. 5E). This was further validated by flow cytometry, where macrophages were identified as the primary producers of CXCL9 in the PAT (Figures. 5F,G), with increased expression following ICB treatment (Figure. 5H). Together, ICB treatment altered the inflammatory microenvironment of the PAT to promote CXCL9 expression in macrophages within the TLS.

**Figure 5.**
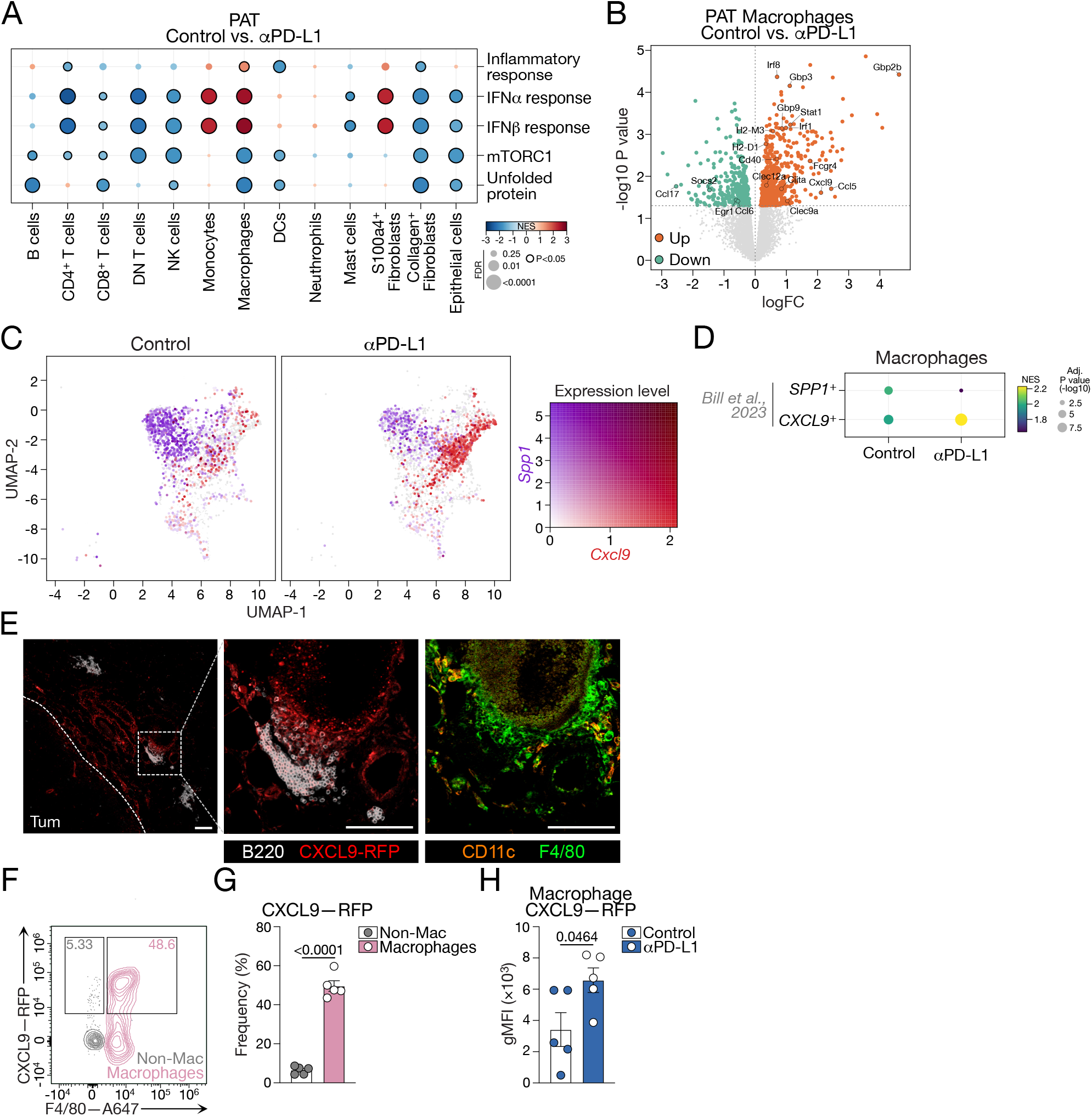
Immune checkpoint blockade induces IFN response in TLS forming niches. (A) Hallmark pathway enrichment across cell types in PAT (anti-PD-L1 vs control). The bubble plot shows GSEA results for five Hallmark pathways in each cell type within PAT. *n* = 4 per group. Positive NES values and red coloring consistently indicate enrichment in anti-PD-L1. Color represents the NES, displayed on a diverging scale symmetrically truncated at ±3, and dot size represents −log10(FDR). Dots outlined in black indicate pathways meeting both P < 0.05 and FDR < 0.25, whereas pale dots without outlines indicate non-significant enrichment. DNT: double negative T; NK: natural killer; DC: dendritic cell. (B) Differential expression in PAT macrophages (anti-PD-L1 vs control). Volcano plot was performed using the edgeR quasi-likelihood framework. The x-axis represents log2 fold change (logFC), with positive values indicating higher expression in anti-PD-L1, and the y-axis represents −log10(*P*). Colored points highlight selected genes of interest, whereas all other genes are shown in grey. (C) *Cxcl9* and *Spp1* gene expression in PAT macrophages. UMAP showing *Cxcl9* and *Spp1* expression in all macrophages from PAT, with control shown on the left and anti-PD-L1 on the right. Both panels use the same coordinate range. Light-grey points show all macrophages as a common background, whereas colored points represent macrophages from the corresponding treatment group. Gene expression is visualized using a two-channel overlay, with red indicating *Cxcl9* and blue indicating *Spp1*. The channels are combined on a white background such that cells co-expressing both genes appear dark purple to near black. For the two-dimensional color key shown at the lower right, expression of each gene was independently normalized to its 99.5th percentile. (D) NES plot of enrichment of human cancer *SPP1* or *CXCL9* expressing macrophage gene signatures. (E) Representative confocal images of TLS in REX3 mice treated with anti-PD-L1. Scale bars represent 100 µm. *n* = 4. White dashed lines represent tumor borders. Tum indicates tumor region. (F) Representative flow cytometry plots for macrophage and nonmacrophage in PAT. (G) Percentage of CXCL9-RFP positive cells for macrophage and nonmacrophage in PAT in (F). *n* = 5. (H) gMFI of CXCL9-RFP in PAT macrophage in control or anti-PD-L1 groups analyzed by flow cytometry. *n* = 5 per group. Data are mean ± SEM. Statistics for (G) was determined using paired *t*-test. Statistics for (H) was determined using unpaired *t*-test.

CXCL9 expression within the tumor microenvironment has been proposed as a biomarker of effective ICB treatment responses and an indicator of favorable tumor outcomes.^62^ The development of the MC38-OVA^lo^ tumor model, enabled us to gain mechanistic understanding of how CXCL9 expression impacts TLS formation and function. When MC38-OVA^lo^ tumors were established in either WT or *Cxcl9^-/-^* hosts, no overt difference was observed in tumor growth (Figure. 6A). However, in contrast to WT mice, tumors in *Cxcl9^-/-^*mice were not responsive to anti-PD-L1 ICB (Figure. 6B). Flow cytometry analysis of PAT indicated decreased immune activity in *Cxcl9^-/-^*mice, with no recruitment of B cells, but similar trends of increases in CD4^+^ and CD8^+^ T cells were observed with ICB (Figure. 6C). However, correlating with the B cell infiltration, proportions of T cell subsets were affected by the loss of CXCL9, where T_FH_ and T_CM_ cells were reduced and T_EX_ cells were increased (Figure. 6D). Additionally, TLS formation was impeded in *Cxcl9^-/-^*hosts, resulting in disorganized immune cell aggregates, rather than organized TLS with segregated T cell and B cell zones (Figure. 6E). Strikingly, ICB did not induce an increase in TLS formation in *Cxcl9^-/-^* mice, as occurred in WT hosts (Figure. 6F). Thus, CXCL9 plays an essential role in recruiting B cells and establishing cellular communication and organization that is fundamental to TLS formation. Lack of CXCL9 reduces the efficacy of ICB, which correlates with decreased TLS-associated T_FH_ and T_CM_ cells and TLS organization.

**Figure 6.**
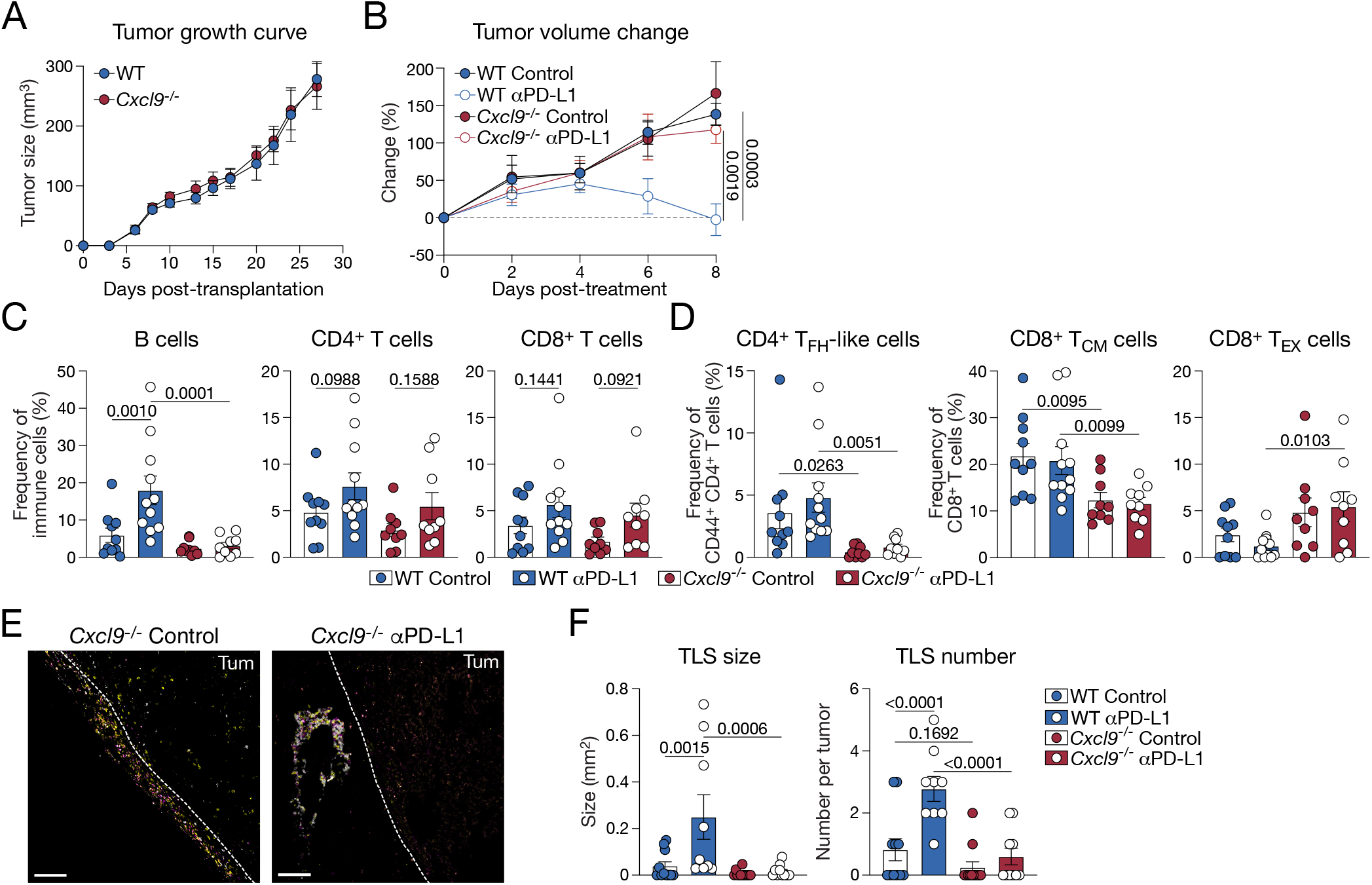
TLS and B cell recruitment is CXCL9 dependent. (A,B) Tumor growth curves and relative tumor size change since anti-PD-L1 treatment for WT and *Cxcl9^-/-^*mice. *n* = 9-11 per group. (C) Frequencies of B cells, CD4^+^ T cells, CD8^+^ T cell in PAT analyzed by flow cytometry. *n* = 9-11 per group. (D) Frequencies of T_FH_-like, CD8^+^ T_CM_, CD8^+^ T_EX_ in PAT analyzed by flow cytometry. *n* = 9-11 per group. T_FH_: T follicular helper; T_CM_: T central memory; T_EX_: T exhausted. (E) Representative confocal images of control or anti-PD-L1 treated MC38-OVA^lo^ peritumoral TLS in *Cxcl9^-/-^*mice. White dashed lines represent tumor borders. Tum indicates tumor region. Scale bars represent 100 µm. (F) Quantifications of TLS size and number in (E). *n* = 9-12 per group. Data are mean ± SEM. Statistics for (A) was determined using unpaired *t*-test. Statistics for (B), (C), (D) and (F) was determined using two-way ANOVA test.

### CXCL9-dependent promotion of antigen-specific responses

Finally, we sought to investigate if the addition of OVA in the MC38-OVA^lo^ model led to antigen-specific immune responses against OVA, and if this correlated with TLS formation and ICB responsiveness. Using single-cell TCR-sequencing (scTCR-seq) analysis, OVA-specific T cells were identified by exact amino-acid matching of TCRβ CDR3 sequences against a curated IEDB ovalbumin reference.^63^ While the number of clones assessed was too low to reach significance, we observed notable infiltration of OVA-specific CD8^+^ T cells in the WT control and anti-PD-L1 treated tumor cores, which were absent in *Cxcl9^-/-^* hosts (Figures. 7A,B). Further, recruitment of OVA-specific T cells to the peritumoral TLS was only observed in WT hosts with anti-PD-L1 treatment (Figures. 7A,B). To assess humoral responses, we first examined GC B cells in the tumor dLN and PAT. Neither proportion of GC B cells differed between WT and *Cxcl9^-/-^* hosts or following anti-PD-L1 treatment (Figure. S5A). Consistent with these findings, the treatment did not increase the tumor specific serum IgG levels or show evidence of clonal selection, as assessed by BCR diversity (Figures. S5B,C). However, we did observe an increase in the tumor specific serum IgM levels in WT anti-PD-L1 treated hosts, that did not occur in *Cxcl9^-/-^* hosts (Figure. 7C,D), suggesting an extrafollicular response that is consistent with IgM production of immature TLS.^5,64^ Additionally, by performing a serum binding assay against all three MC38 cell lines, we observed that antibody responses were not specifically directed towards the OVA model antigen, as binding was comparable irrespective of the level of OVA expression (Figure. 7E). Thus, while there is some evidence of CXCL9-dependent antigen-specific responses, we suggest that TLS B cells may play a multifaceted role in anti-tumor immunity, that could include structural formation of TLS, antigen presentation and IgM antibody production.

**Figure 7.**
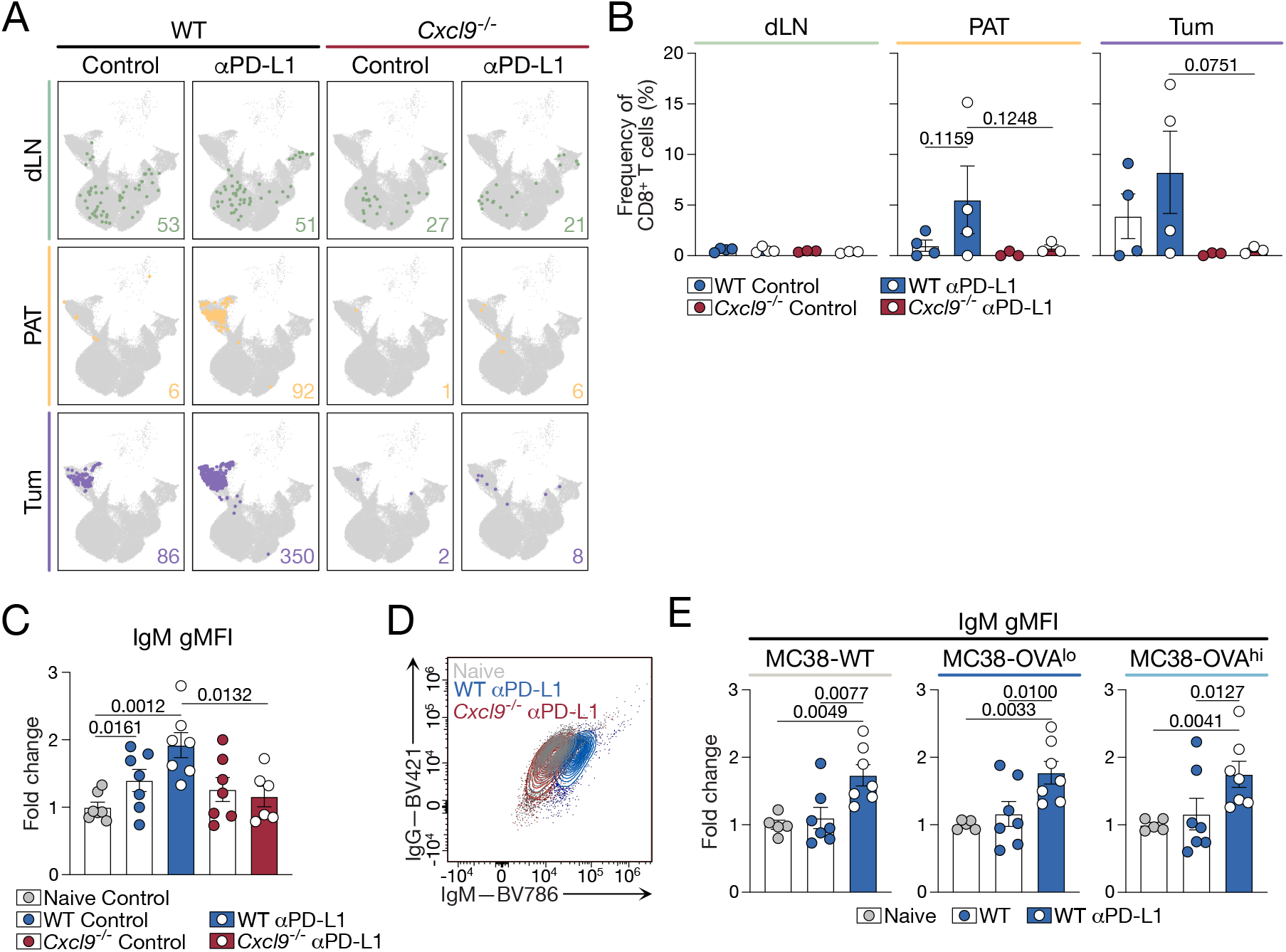
Immune checkpoint blockade promotes antigen-specific responses in CXCL9-dependent manner. (A) Distribution of OVA-specific T cells across tissues and treatment groups. OVA-specific T cells were identified by exact amino-acid matching of TCRβ CDR3 sequences against a curated IEDB ovalbumin reference. Matching was performed using the IMGT-defined CDR3 core, with the conserved N-terminal C and C-terminal F/W residues removed. Each panel shows the same UMAP embedding of all 42,440 T cells in grey, with OVA-specific T cells highlighted. The number of OVA-specific T cells in each subset is indicated in the lower-right corner of each panel. (B) Per-mouse frequency of OVA-specific T cells across tissues and treatment groups in (A). *n* = 3-4 per group. (C) gMFI fold change of serum IgM binding to MC38-OVA^lo^ cells, relative to naïve serum analyzed by flow cytometry. *n* = 6-7 per group. (D) Representative flow cytometry plots for naïve, WT anti-PD-L1 and *Cxcl9^-/-^*anti-PD-L1 serum antibody binding to MC38-OVA^lo^ cells in (C). (E) gMFI fold change of serum IgM binding to MC38-WT, MC38-OVA^lo^ and MC38-OVA^hi^ cells, relative to naïve serum analyzed by flow cytometry. *n* = 5-7 per group. Data are mean ± SEM. Statistics for (B), (C) and (E) was determined using two-way ANOVA test.

## DISCUSSION

The lack of mouse models that faithfully recapitulate human TLS development has limited mechanistic investigation of how these structures form and function. Here, we have established and characterized a subcutaneous MC38 tumor model that spontaneously generates organized peritumoral TLS. Using confocal imaging, flow cytometry, and scRNA-seq, we demonstrate that MC38-OVA^lo^ tumors develop TLS with cellular and structural features resembling immature human TLS. This model extends existing preclinical systems by providing a tractable platform to investigate TLS formation, maturation, and their modulation by immunotherapies such as ICB. Our findings further identify the IFN-inducible chemokine CXCL9 as a functional link between the inflammatory tumor niche, TLS formation, and effective ICB responses.

Comparison of MC38 tumor variants suggest that TLS formation is influenced not simply by tumor antigen expression, but by the kinetics and persistence of tumor-immune engagement. MC38-OVA^hi^ tumors can be cleared, whereas MC38-WT tumors escape immune control to develop an overwhelming tumor burden. In contrast, MC38-OVA^lo^ tumors occupy a “Goldilocks” range of antigenicity, where expression is sufficient to engage anti-tumor immunity, but not so high that the tumor is eliminated. This intermediate state supports slow, sustained tumor growth and prolonged interaction between the tumor and immune cells. Importantly, serum IgM antibodies generated in mice bearing MC38-OVA^lo^ tumors bound the parent MC38-WT cells at comparable levels, indicating that the humoral response recognized antigens shared by the two tumor lines. Thus, OVA expression is not the sole driver of tumor antigen recognition. We therefore propose that, rather than a single initiation event, sustained immune engagement generated by slow tumor growth remodels the tumor border and creates conditions that favor TLS formation. Preservation of the peritumoral stroma and adipose tissue is technically challenging during tumor excision, which may help explain why TLS present in this compartment have been overlooked in subcutaneous tumor models.

The anatomical context of TLS formation may be as important as tumor antigenicity. In human cancers, TLS can develop within the tumor core or in adjacent peritumoral tissues. The peritumoral region serves as a critical interface where immune cell infiltration, vascular organization, and stromal signals create a niche permissive for TLS development and maturation.^31,39,40,42,65^ Several reported mouse models with spontaneous TLS formation are based on lung cancer.^32,45,46^ This may reflect the intrinsic capacity of the lung to organize ectopic lymphoid structures following respiratory infection and during autoimmune diseases.^66^ Our findings extend this concept to the subcutaneous compartment, showing that peritumoral adipose tissue can also provide a permissive niche for TLS formation, when sustained tumor-immune engagement is present. Within this niche, spatially restricted expression of CCL21 and CXCL13 demarcated T cell and B cell zones, indicating that chemokine organization is a defining feature of TLS architecture.

TLS appear to function as an intermediate lymphoid site between the tumor draining lymph node and the tumor. Within the peritumoral TLS, we identified a population of TCF-1^+^ T_CM_ cells. This population resembles T_SL_ cells that are maintained in chemokine-defined niches within lymph nodes, where they support the ongoing generation of effector cells.^67–70^ Their presence within TLS suggests that these structures may provide a local reservoir of tumor-reactive T_SL_ cells and a site for replenishing effector or T_EX_ cell populations within the tumor.^20^ The B cells were similarly spatially organized, with high proportions of IgD^+^ naïve-like B cells within the peritumoral TLS, while activated IgD^-/low^ B cells were predominant inside the tumor border. Similarly, we found multiple lymphoid cell states at frequencies that bridge lymph node and tumor populations, consistent with the notion that TLS act as an interface between these tissues.

Using the ICB-sensitive MC38-OVA^lo^ model, we demonstrated that anti-PD-L1 promoted TLS formation and expansion. This was associated with increased B cell recruitment and immune activity within the peritumoral niche rather than the tumor core. These observations provide mechanistic insight as to why TLS are a predictive biomarker of favorable responses to ICB across a range of human cancers.^6,7,51^ However, the role of B cells in ICB responsiveness and cancer prognosis in general has not been clearly elucidated.^64,71^ Naïve-like B cells in TLS express higher levels of chemokine receptors and costimulatory markers than naïve B cells in the lymph nodes, suggesting that they may be functionally poised to present antigen and support T cell activation. Although CXCL9-dependent B cell recruitment was required for TLS formation and ICB responses, these B cells did not express the receptor CXCR3 (not shown). Thus, CXCL9 may act indirectly as part of an inflammatory cascade with other chemoattractants, such as EBI2 and CXCL12, to recruit naïve-like B cells to TLS.^31,72^ Additionally, we observed reduced *Dpp4* interactions in peritumoral tissue following ICB. DPP4 is an enzyme that cleaves chemokines, therefore reduced DPP4 activity may preserve local chemokine gradients to promote cell interaction, recruitment and expansion.^52^ It is undetermined if this is conserved in human TLS, however human B cells have been reported to increase DPP4 expression upon activation.^73^ Despite TLS expansion following ICB, the anti-tumor response was dominated by IgM, rather than IgG. Together with the absence of major changes in GC B cells or BCR clonal selection, this finding suggests that tumor-specific B cells in the lymph nodes or TLS undergo extrafollicular differentiation rather than classical GC responses.^64^ Combined, in this model, tumor-specific B cells may not only contribute to the structural formation of TLS, but also to T cell costimulation, chemokine retention and antibody production.

CXCL9 expression is associated with immune-hot niches within tumors and correlates with effective ICB responses.^18,19,35^ Our findings further identify CXCL9 as a necessary regulator of both TLS formation and expansion and ICB efficacy. In the absence of CXCL9, this spatial immune network is blocked and could not be rescued by anti-PD-L1. *Cxcl9^-/-^* mice failed to develop baseline TLS, yet exhibited comparable tumor growth to wildtype hosts. This suggests that immature TLS are insufficient to constrain tumor growth and that ICB-dependent TLS functionality is required for effective tumor control. Although *Cxcl9^-/-^* mice maintained a similar proportion of CD8^+^ T cells as WT hosts, they failed to expand or recruit OVA-specific CD8^+^ T cells into the peritumoral tissue and tumor core. Furthermore, the T cells that were present in *Cxcl9^-/-^* hosts displayed a less TLS-associated phenotype, characterized by reduced T_FH_ cell and T_CM_ cell populations and an accumulation of T_EX_ cells. Together, CXCL9 supports both recruitment of tumor antigen-specific T cells and catalyzes the cell interactions that organize functional TLS.

We found that CXCL9-expressing macrophages populated TLS and interacted with B cells following ICB treatment, positioning macrophage-derived CXCL9 as a potential link between inflammation, immune cell recruitment, and TLS organization. These findings raise the possibility that selectively enhancing macrophage-derived CXCL9 while preserving the appropriate cellular composition of the niche may provide a therapeutic route to promote TLS formation and improve responses to ICB.

Together, our findings suggest that TLS formation is not simply a consequence of high tumor immunogenicity. Rather, TLS emerge when an intermediate level of tumor antigenicity generates sustained tumor-immune engagement within a tissue niche capable of supporting organized immune interactions. The MC38-OVA^lo^ model provides a tractable system for studying this process and for testing strategies to induce, remodel and therapeutically exploit TLS in solid tumors.

### Experimental models

#### Animals

Mice were maintained on a C57BL/6J background in specific pathogen-free conditions. Reporting the Expression of CXCR3 ligands (REX3) transgenic,^61^ *Cxcl9^−/−^*,^74^ and Nur77-GFP mice,^75^ have been described previously. Mice used in this study were 6-10 weeks of age and sex-matched. Animal experiments were performed in accordance with the Walter and Eliza Hall Institute (WEHI) animal ethics committee.

#### Tumor model

Ovalbumin (OVA) expressing MC38-OVA^lo^ and MC38-OVA^hi^ colon carcinoma cell lines were generated by transducing MC38 cell line with pFUGW-OVAC1 lentiviral vectors. Sorted single cells with low and high expression of OVA were then clonally expanded. All cell lines were cultured in Dulbecco’s Modified Eagle Medium (ThermoFisher Scientific) with 10% heat inactivated fetal calf serum in 10% CO_2_ and 37°C incubators. Cells were split and medium was refreshed every 2-3 days.

For tumor transplantation, 10^6^ MC38, MC38-OVA^lo^ or MC38-OVA^hi^ cells were injected subcutaneously into the left flank of mice. Tumor size was measured every 2-3 days with an electronic caliper and reported as tumor size using formula (tumor length)×(tumor width)^2^’p’(4/3).

### Method details

#### Treatment of tumor-bearing mice

For anti-PD-L1 immune checkpoint blockade therapy, on day 22, 25 and 28 post tumor transplantation, tumor-bearing mice were treated intraperitoneally with 200μg anti-mouse PD-L1 antibodies (B7-H1, BioXcell) or rat IgG control (WEHI antibody facility).

#### Tumor tissue preparation

To preserve the tertiary lymphoid structure (TLS) integrity, tumors were harvested with peritumoral tissues. For imaging, the skin, stroma/adipose tissue and peritoneal membrane attached to the tumor were removed as a whole with the tumor into fixative solution.

For flow cytometry and scRNA-seq, peritumoral stroma/adipose tissues were removed with the tumor. Peritumoral tissues were then macroscopically separated from tumor core by peeling them off from the tumor capsule with forceps under a dissecting microscope. Tumor cores and peritumoral tissues were mechanically disrupted with scissors and incubated in RPMI-1640 with 20μg/ml DNase I (Roche) and 1mg/ml Collagenase IV (Sigma-Aldrich) for 30 minutes at 37°C. Cells were filtered through a 70μm cell strainer (Falcon) to achieve single cell suspension. Tumor core cells were added carefully on top of Ficoll-Paque (GE Health Care). Ficoll gradient was spun at 2000rpm for 30 minutes at room temperature with zero-level acceleration and zero-level break. The layer containing immune cells was collected and washed in FACS buffer.

#### Flow cytometry

Single cell suspensions of draining lymph nodes, peritumoral tissue and tumor core were incubated with FcBlock (BD Biosciences) for 10 minutes at room temperature. Then cells were stained with CXCR5 antibody for 1 hour at 37°C, followed by all other surface staining antibodies for 30 minutes at 4°C. After washing, cells were stained with viability dye staining for 10 minutes at 4°C. Transcription factor staining was performed using Foxp3 Transcription Factor Staining Kit (Thermo Fisher Scientific). All flow cytometry analysis was performed on BD FACSymphony A3 Cell Analyzer (BD Biosciences) and Cytek Aurora (Cytek Biosciences). Data analysis was performed with FlowJo v.10 (FlowJo LLC).

#### Detection of plasma immunoglobin binding to tumor cells

Cultured MC38-WT/MC38-OVA^lo^/MC38-OVA^hi^ cells were dissociated by Versene (Gibco). Cells were stained with viability dye for 10 minutes at 4°C and blocked with FcBlock (BD Biosciences) for 10 minutes at 4°C. Mouse blood serum was collected from respective tumor bearing mice and added to reach 1:10 dilution and incubated for 1 hour at 4°C. Cells were washed with FACS buffer and incubated with 1:100 diluted rat anti-mouse IgM BV786 (II/41) and 1:100 diluted goat anti-mouse IgG BV421 for 30 minutes at 4°C. Cells were washed, fixed and washed again for flow cytometry analysis on BD FACSymphony A3 Cell Analyzer (BD Biosciences) to quantitatively determine the mouse IgM and IgG binding to the tumor cell surface.

#### Immunofluorescence and confocal microscopy

Tumors were fixed in 4% paraformaldehyde (PFA) at 4°C for 24 hours and immersed in 30% sucrose for an additional 24 hours at 4°C before being embedded in optimal cryostat temperature (OCT). Across the whole tumor mass, tissue sections were acquired at several positions, which were 200-300μm apart. For each position, three to four 20μm serial sections were cut via microtome onto SuperFrost Plus slides.

Sections were blocked in PBS containing 0.1% Triton X-100 (Sigma-Aldrich) and 1% BSA (Sigma-Aldrich) for 30 minutes at room temperature and stained overnight at 4°C in PBS containing 0.01% Triton X-100 and 0.1% BSA with primary antibodies. Where applicable, slides were washed twice and stained with secondary antibodies for 1 hour at room temperature. Slides were washed in 0.1% Triton X-100 twice and in PBS twice and coverslips were mounted with ProLong Gold (Thermo Fisher Scientific). Images were acquired using a LSM980 confocal microscope using 20× or 63× objective (Carl Zeiss MicroImaging). The acquisition software was Zen Blue (ZEISS).

#### Definition and quantification of TLS

Peritumoral TLSs were identified by tight aggregates of 50 or more B cells in juxtaposition to PNAd^+^ vasculature.^65,76^ TLS areas were quantified by polygon tracing (Zen Blue). TLS numbers per tumor were determined by counting TLS in tissue sections. Boundaries of the tumor were defined by tissue morphology.

#### Sample preparation for scRNA-seq

After acquiring single suspensions following tissue preparation, dLN cells were stained with fluorescent antibodies and Hashtags for 30 minutes at 4°C, followed by viability staining for 10 minutes at 4°C. Peritumoral and tumor single cells were stained with Hashtag separately, followed by viability staining for 10 minutes at 4°C. Sample suspensions were sorted using BD FACSAria Fusion Flow Cytometer (BD Biosciences) to isolate live cells from peritumoral and tumor samples and 4 cell populations from draining lymph node samples for scRNA-seq and single cell T cell receptor and B cell receptor sequencing (scTCR/BCRseq).

#### scRNA-seq data analysis

Single-cell gene expression data from Cell Ranger (v9) output was analyzed using standard Scanpy (v.1.9.3) workflow.^77,78^ Cell barcodes were demultiplexed with Hashtag oligos and barcodes flagged as multiplet, negative or uncertain were removed. Cell Ranger and hashtag demultiplexing were performed by the WEHI SCORE team. Further downstream preprocessing included doublet detection using Scrublet (v0.2.3)^79^ with adaptations outlined by Popescu et al.^80^ Briefly, after scrublet was performed, the data was iteratively sub-clustered and a median scrublet score for each sub-cluster was computed. Median absolute deviation (MAD) scores were computed from the cluster scrublet scores and a one tailed t-test was performed with Benjamini-Hochberg (BH) correction^81^ applied and cells with significantly outlying cluster scrublet scores (BH P < 0.1) were flagged as potential doublets. The data was then processed using Scanpy with standard quality control steps; cells were filtered if number of genes > 6000 or < 200. Mitochondrial content cut-off was set at 30%. Genes were retained if they are expressed by at least 3 cells. Genes counts for each cell were normalized to contain a total count equal to 10000 counts per cell. This led to a working dataset of 10,288 cells. Highly variable genes were selected based on the following parameters: minimum and maximum mean expression are >=0.0125 and <=3 respectively; minimum dispersion of genes = 0.5. The number of principal components used for neighborhood graph construction and dimensional reduction was set at 40. Clustering was performed using Leiden algorithm^82^ with resolution set at 1.0. Size of local neighborhoods was set at 10. Uniform Manifold Approximation and Projection (UMAP; v0.5.7)^83^ was used for dimensional reduction and visualization, the minimum distance was set at 0.3 and all other parameters as per default settings in Scanpy. Cell type identification was guided by manual inspection of canonical marker gene expression and top marker genes identified with Wilcoxon rank sum tests.

#### Sample processing, single-cell analysis and cell-type annotation

Tum, PAT and dLN samples from 14 mice were analyzed across four experimental groups: WT Ctrl, WT anti-PD-L1, *Cxcl9^-/-^* Ctrl and *Cxcl9^-/-^* anti-PD-L1. Samples were multiplexed using hashtag oligonucleotides (HTOs), yielding 42 biological samples together with an HT-2 hashing control, distributed across four 10x Genomics capture batches. Sequencing data were aligned to the mouse GRCm39 reference genome using Cell Ranger, and samples were demultiplexed according to HTO signals using demuxmix.^84^ Cells were filtered based on library quality-control metrics, and potential doublets were identified using Scrublet.^79^ The final single-cell transcriptomic dataset contained 140,770 cells and 33,696 genes.

Gene expression was library-size normalized to 10,000 counts per cell followed by log1p transformation. A total of 2,083 highly variable genes were selected using the Seurat method implemented in Scanpy,^77^ followed by z-score scaling and principal component analysis (PCA). UMAP embedding was then performed, and cells were clustered using the Leiden algorithm, resulting in 33 clusters. Cell types were manually annotated based on the expression of canonical marker genes.

#### Pseudobulk differential expression analysis

Differential expression analysis was performed at the pseudobulk level, with individual mice treated as biological replicates. Differential expression was assessed using edgeR.^85^ Lowly expressed genes were filtered using filterByExpr function in edgeR, and library composition was normalized using the trimmed mean of M-values (TMM) method.^86^ Dispersion was estimated using estimateDisp, followed by quasi-likelihood model fitting and testing with glmQLFit and glmQLFTest. P values were adjusted for multiple testing using the Benjamini–Hochberg procedure.

#### CD8^+^ T cell differentiation trajectory analysis

Trajectory analysis was performed on CD8^+^ T cells carrying TCR clonotype information, comprising 22,760 cells in total. An ElPiGraph elastic principal curve was fitted in the CD8^+^ T cell UMAP space using scFates with 30 principal points, mu = 0.1 and lambda = 0.01.^87,88^ The trajectory was oriented from the naïve CD8^+^ T cell-enriched region towards the CD8^+^ T_EX_-enriched region. To avoid non-biological detours caused by isolated cell islands in the UMAP embedding, the principal curve was fitted only to the largest connected component of the UMAP.

#### TCR clonotype similarity clusters

TCR clonotype similarity clusters were defined from the tcrdist-derived cc_aa_tcrdist assignments.^89^ The cc = 20 cluster shown in Figure. 3G was the largest CD8^+^ T cell clonotype similarity cluster and contained 1,127 cells. This cluster comprised 1,012 distinct TCR clonotypes and therefore represents multiple clonotypes with similar TCR sequence features rather than a single expanded clone.

#### Cell-cell interaction analysis

Cell-cell interaction analysis was performed using the raw expression matrix from the annotated single-cell object. The matrix contained 33,696 genes with non-negative log1p-normalized expression values. HT-2 spike control samples and cells annotated as Multiplet were excluded prior to analysis. Mouse gene symbols were mapped to human orthologues using mousipy. Statistical analysis method was performed independently for each biological sample. For each ligand–receptor and cell-type pair, interaction strength was summarised by averaging CellPhoneDB means across mice within each condition.^90^

#### Gene Set Enrichment Analysis (GSEA) analysis

Preranked gene-set analysis on mouse MSigDB Hallmark gene sets was performed using preranked gene lists with the fgsea package in R.^91–93^ Genes were preranked according to pseudobulked differential expression gene testing results from edgeR (v4.4.2) (for each cell-type, group and tissue separately; Log-foldchange-signed −log10 P values).^94^

#### Antigen specific TCR analysis

Paired αβ TCR sequencing data were processed using Dandelion (v0.5.0).^95^ Following per-cell filtering, the dataset contained 79,341 productive TCR contigs from 43,951 cells, of which 42,395 cells carried a single productive TCRβ contig. A reference set of OVA-reactive TCRs was obtained from the IEDB database.^63^ Only receptor entries for which the epitope source molecule contained “ovalbumin” were retained, yielding 383 reference receptors, of which 364 were H2-Kb restricted. CDR3 amino-acid sequences were further converted to IMGT-defined CDR3 cores by removing the conserved N-terminal cysteine and the C-terminal phenylalanine or tryptophan. This yielded 331 unique TCRβ CDR3 core sequences but only two TCRα CDR3 cores; therefore, subsequent matching was performed using the β chain alone. A T cell was defined as OVA-associated when its TCRβ CDR3 core exactly matched, at the amino-acid level, any β-chain CDR3 core in the reference set. The frequency of OVA-associated T cells was calculated as their percentage among all T cells within the corresponding comparison unit. All analyses were performed using scanpy v1.11.0, pandas v2.3.3, and seaborn v0.13.2. UMAP coordinates and cell-type annotations were obtained directly from the previously annotated transcriptomic object and were not recomputed.

#### BCR α diversity analysis

BCR clones were assigned using Dandelion’s heavy-chain clone definition. Only B cells carrying a productive IGH sequence with a valid clone assignment were retained. For cells with multiple productive IGH sequences, the sequence with the highest UMI count was selected. Clonal abundance was determined by cell counts, with each cell weighted equally.

BCR clonal diversity was compared across the four treatment groups within each tissue using alakazam v1.2.1.^96^ Groups with at least 10 B cells were included, and treatment groups within each tissue were resampled to the cell count of the smallest group to control for differences in sampling depth. Diversity was calculated using Hill numbers over q = 0–4 in increments of 0.1, with q = 0, 1 and 2 corresponding to clonal richness, exponential Shannon diversity and inverse Simpson diversity, respectively. Diversity estimates and 95% confidence intervals were obtained from 200 bootstrap resamples. Lines and shaded regions represent the mean bootstrap estimates and 95% confidence intervals, respectively. Group differences were assessed using the bootstrap delta test.

#### Human correlation analysis

Enrichment of marker gene sets derived from published datasets (Figures. 2J,3I,5D)^6,22,60,64,71,97^ was assessed using the *fgsea* package.^98^ For Figures 2J and 5D, genes were ranked according to their mean expression across cells within each indicated group (dLN and PAT for naïve B cells in Figure 2J; Control and αPD-L1 for macrophages in Figure 5D). Ranked gene lists were used as input for *fgsea* to calculate normalized enrichment scores (NES) for the published marker gene sets. Results were visualized as dot plots, with dot size representing −log10-transformed adjusted *P* values and dot color representing NES. For Figure 3I, NES analysis was performed using pseudobulk samples generated by aggregating WT control cells according to sample, cell type, and experimental group, with a minimum of 30 cells per pseudobulk sample. Genes were filtered using *edgeR::filterByExpr* function based on design matrix incorporating these covariates (i.e. cell type, and experimental group). Gene expression counts were normalized using the TMM method. The resulting normalized expression profiles of CD4^+^ and CD8^+^ T cells in the PAT were used to generate ranked gene lists and calculate NES for the relevant marker gene sets using *fgsea*. NES values and statistical significance were visualized as dot plots.

#### Software

Single-cell analyses were primarily performed in Python 3.13 using Scanpy 1.11.0, AnnData 0.11.3, NumPy 2.1.3, SciPy 1.15.2, pandas 2.3.3, scFates and Matplotlib 3.10.3. Pseudobulk differential expression analysis and ortholog mapping were performed in R 4.4.3 using edgeR 4.4.2, limma 3.62.2, and babelgene.

#### Statistics and reproducibility

Statistical differences between groups were evaluated by paired or unpaired Student’s *t*-tests or one-way ANOVA. P values < 0.05 were considered statistically significant. All experimental data are presented as the mean ± standard error of the mean (S.E.M.) with statistical analysis performed using Prism 8 (GraphPad Software). Graphs were generated using Prism 8. Number of biologically independent samples in each group is denoted as “*n*”.

## Data availability

Data are in the article itself and its supplementary materials. scRNA-seq and scTCR/BCR-seq data will be deposited at GEO upon publication. All experimental models and reagents will be made available upon installment of a material transfer agreement.

## Acknowledgements

We acknowledge the Wurundjeri people of the Kulin nation as the traditional owners and guardians of the land on which the majority of the work was performed. We thank Stephen Nutt (WEHI) and Jessica Da Gama Duarte (Monash University) for discussion and feedback on the manuscript. scRNA-seq and scTCR/BCR-seq experiments were technically assisted by the WEHI Genomics Facility. This work was supported by U.S. Department of Defense Melanoma Ideas Award (HT9425-23-1-0967). J.R.G. is an NHMRC Investigator (GNT2007812). R.Z.Q. and B.J.B. were supported by Melbourne Research Scholarship and Tour de Cure PhD Scholarship. B.E.W. is supported by Melbourne Research Scholarship. H.A. is supported by a Walter Benjamin Programme DFG fellowship. This work was made possible through Victorian State Government Operational Infrastructure Support and Australian Government NHMRC IRIISS.

## Author contributions

Conceptualization: R.Z.Q. and J.R.G.

Methodology: R.Z.Q. and G.M.

Investigation: R.Z.Q, H.A., G.M., B.J.B., B.E.W., C.W.T., B.C.D.

Software and Formal Analysis: R.Z.Q., G.M., C.W.T.

Visualization: R.Z.Q., G.M. and B.J.B.

Resources: M.R.J., N.D.G., K.L.R., Z.K.T., J.R.G.

Writing – Original draft: R.Z.Q. and J.R.G.

Funding Acquisition: J.R.G.

Supervision: Z.K.T. and J.R.G.

---

**Figure S1.**
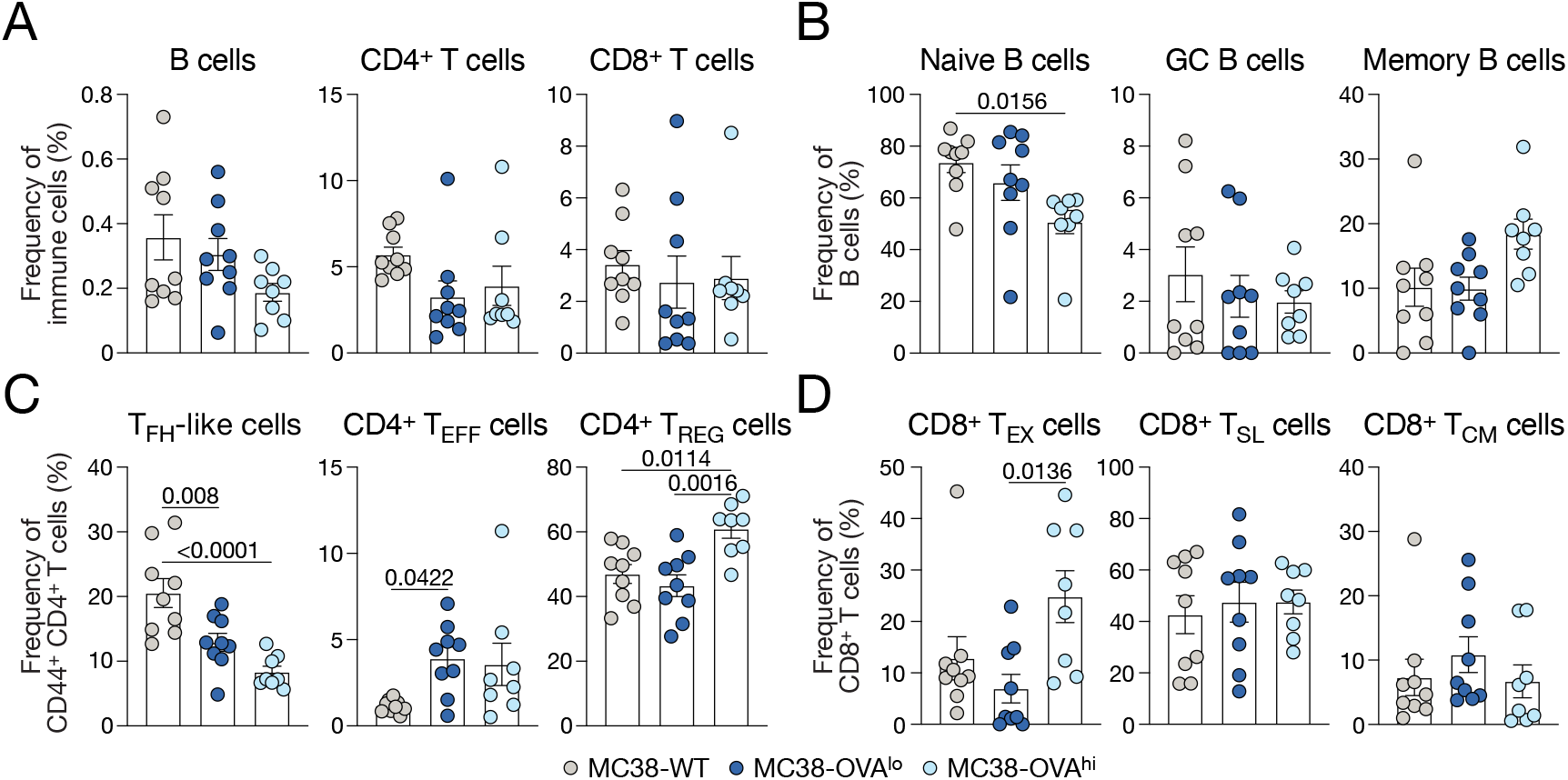
Flow cytometry analysis of tumor associated tissues from MC38-WT, MC38-OVA^lo^ and MC38-OVA^hi^ tumors. (A-D) Frequencies of B and T cell subsets of MC38-WT, MC38-OVA^lo^ and MC38-OVA^hi^ tumors day 15 post transplantation analyzed by flow cytometry. GC: germinal center; T_FH_: T follicular helper; T_EFF_: T effector; T_REG_: regulatory T; T_EX_: T exhausted; T_SL_: stem-like T; T_CM_: T central memory. *n* = 8-9 per group. Data are mean ± SEM. Statistics for (A-D) was determined using one-way ANOVA test.

**Figure S2.**
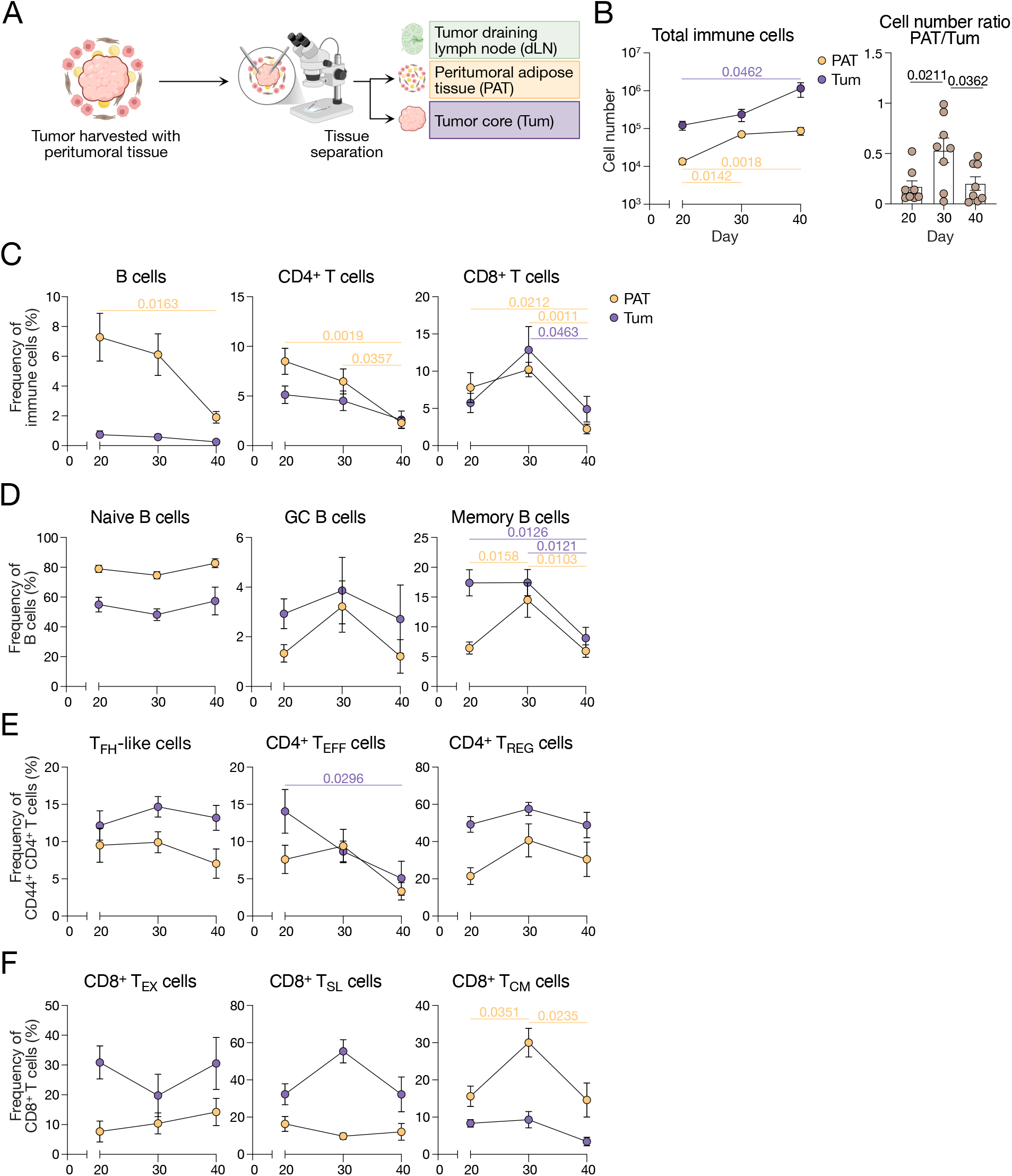
Timecourse analysis of MC38-OVA^lo^ tumors on day 20, 30 and 40 post tumor transplantation. (A) Schematic of tissue processing. (B) Total immune cell number and PAT and Tum cell number ratio across three timepoints post MC38-OVA^lo^ transplantation analyzed by flow cytometry. *n* = 8 per group. (C-F) Frequencies of B and T cell subsets across 20, 30, 40 days post MC38-OVA^lo^ transplantation analyzed by flow cytometry. GC: germinal center; T_FH_: T follicular helper; T_EFF_: T effector; T_REG_: T regulatory; T_EX_: T exhausted; T_SL_: T stem-like; T_CM_: T central memory. *n* = 8 per group. Data are mean ± SEM. Statistics for (B-F) was determined using one-way ANOVA test.

**Figure S3.**
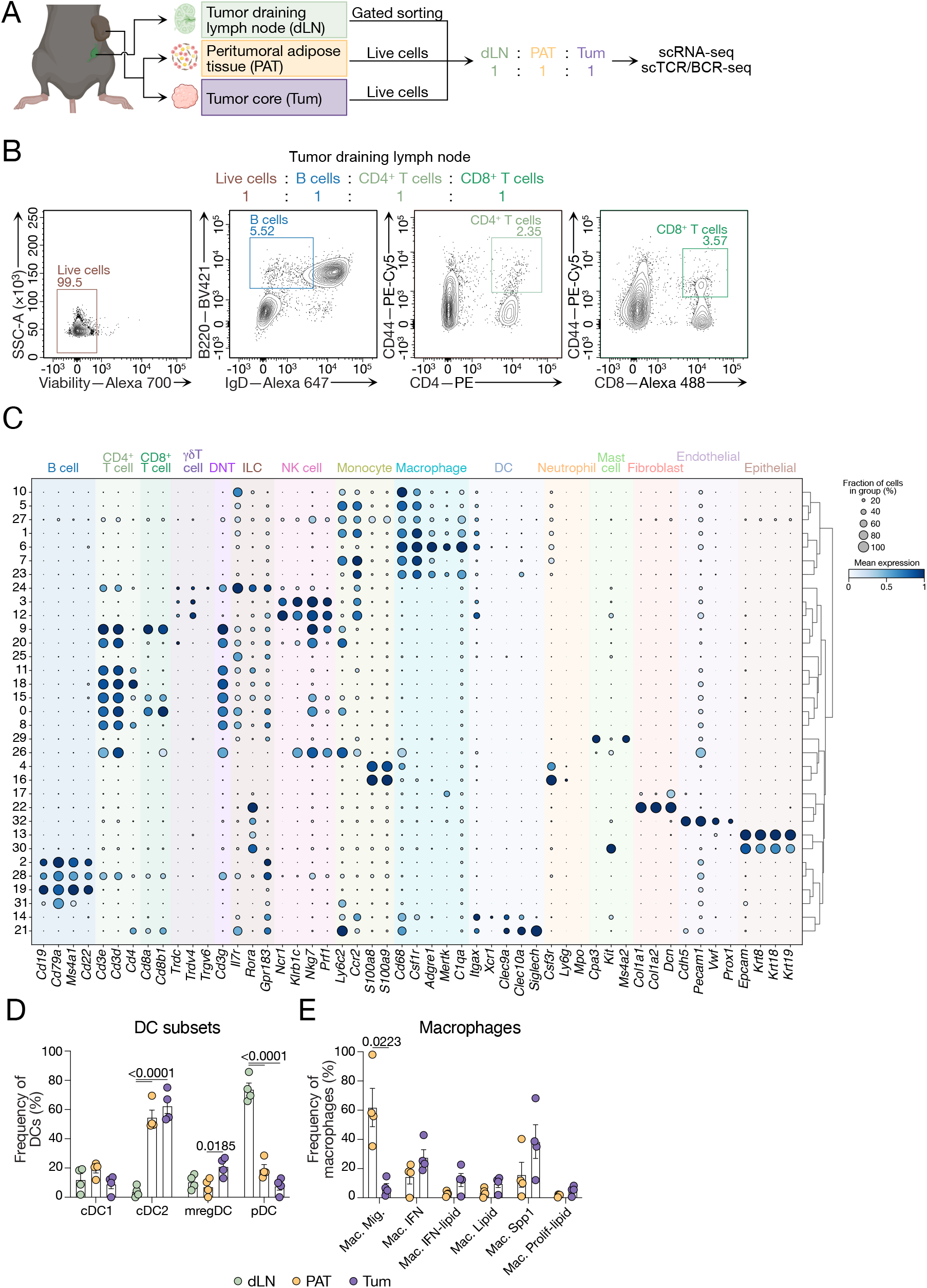
Single cell analysis identifies 15 broad cell types and shows tissue specific myeloid cell subset enrichment. (A) Experimental scheme of scRNA-seq experiment. PAT, Tum and dLN of MC38-OVA^lo^ tumors were harvested on day 30 post transplantation. Live cells in PAT, live cells in Tum and 4 cell populations in dLN were sorted and combined with 1:1:1 ratio prior to sequencing. (B) Gating strategies for 4 cell populations in dLN in (A), i.e. all lymph node lymphocytes (P1), activated B cells (P2), activated CD8^+^ T cells (P3) and activated CD4^+^ T cells (P4). (C) Marker-gene dot plot supporting the cell-type annotations. Rows represent the 33 Leiden clusters across the full dataset, ordered according to hierarchical clustering; columns represent canonical marker genes grouped by the broad cell types they define. Dot size indicates the fraction of cells within each cluster expressing the corresponding gene, while color represents mean expression, min–max scaled to [0, 1] for each gene across clusters. DNT: double negative T cell; NK: natural killer; ILC: innate lymphoid cell; DC: dendritic cell. (D,E) Frequencies of dendritic cell and macrophage subsets from scRNA-seq. cDC1/2: conventional dendritic cell type 1/2; mregDC: mature dendritic cells enriched in regulatory molecules; pDC: plasmacytoid dendritic cell; Mac.: macrophage; Mig.: migratory; IFN: interferon; Prolif: proliferating. *n* = 4. Data are mean ± SEM. Statistics for (D) was determined using paired one-way ANOVA test. Statistics for (E) was determined using paired *t*-test.

**Figure S4.**
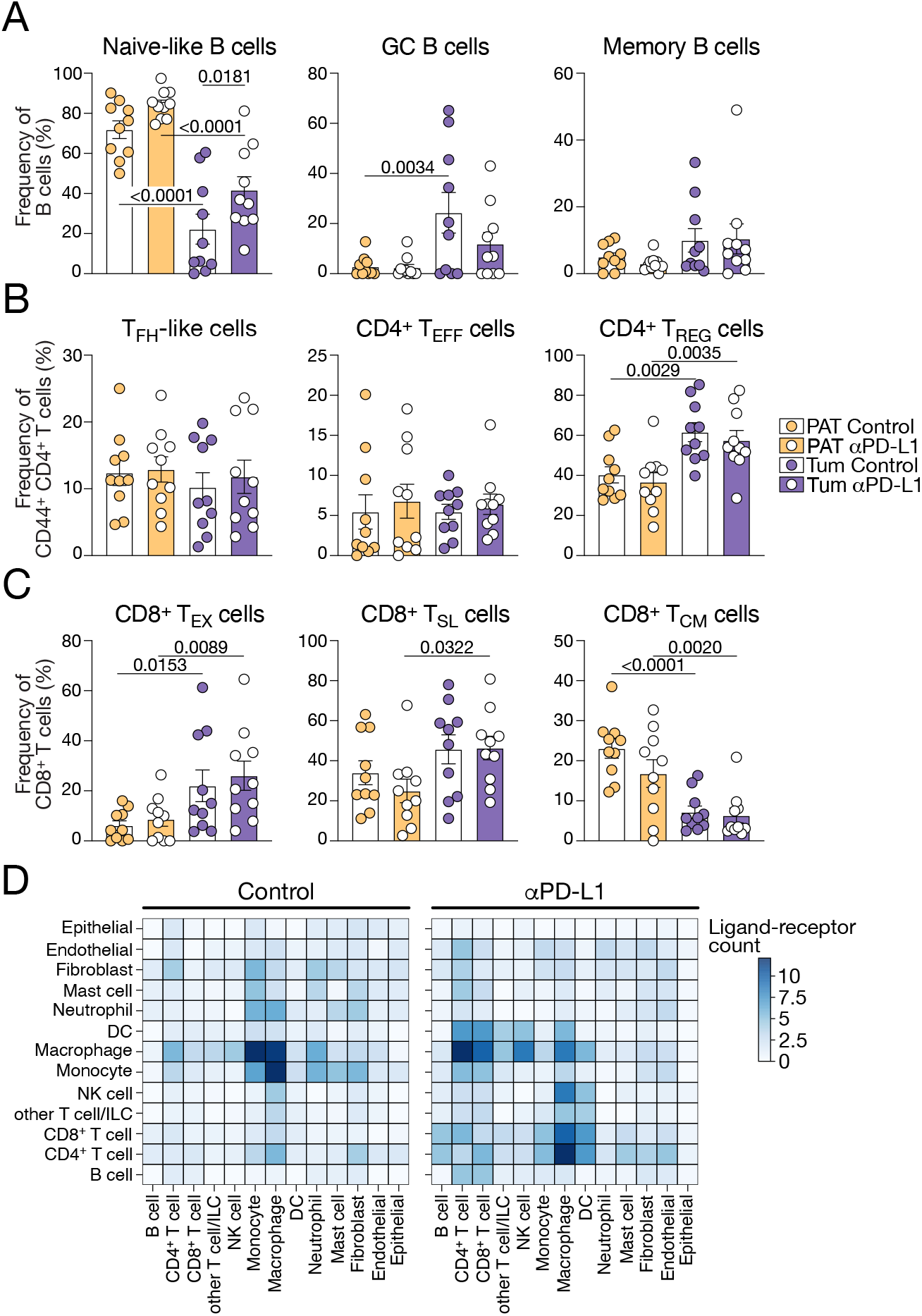
Anti-PD-L1 treatment induces no change in B and T cell phenotypes. (A-C) Frequencies of B and T cell subsets in PAT and Tum for control or anti-PD-L1 treated mice analyzed by flow cytometry. GC: germinal center; T_FH_: T follicular helper; T_EFF_: T effector; T_REG_: T regulatory; T_EX_: T exhausted; T_SL_: T stem-like; T_CM_: T central memory. *n* = 10 per group. (D) Mean number of significant ligand receptor interactions in Tum for control or anti-PD-L1 treated mice, analyzed from scRNA-seq. Data are mean ± SEM. Statistics for (A-C) was determined using two-way ANOVA test.

**Figure S5.**
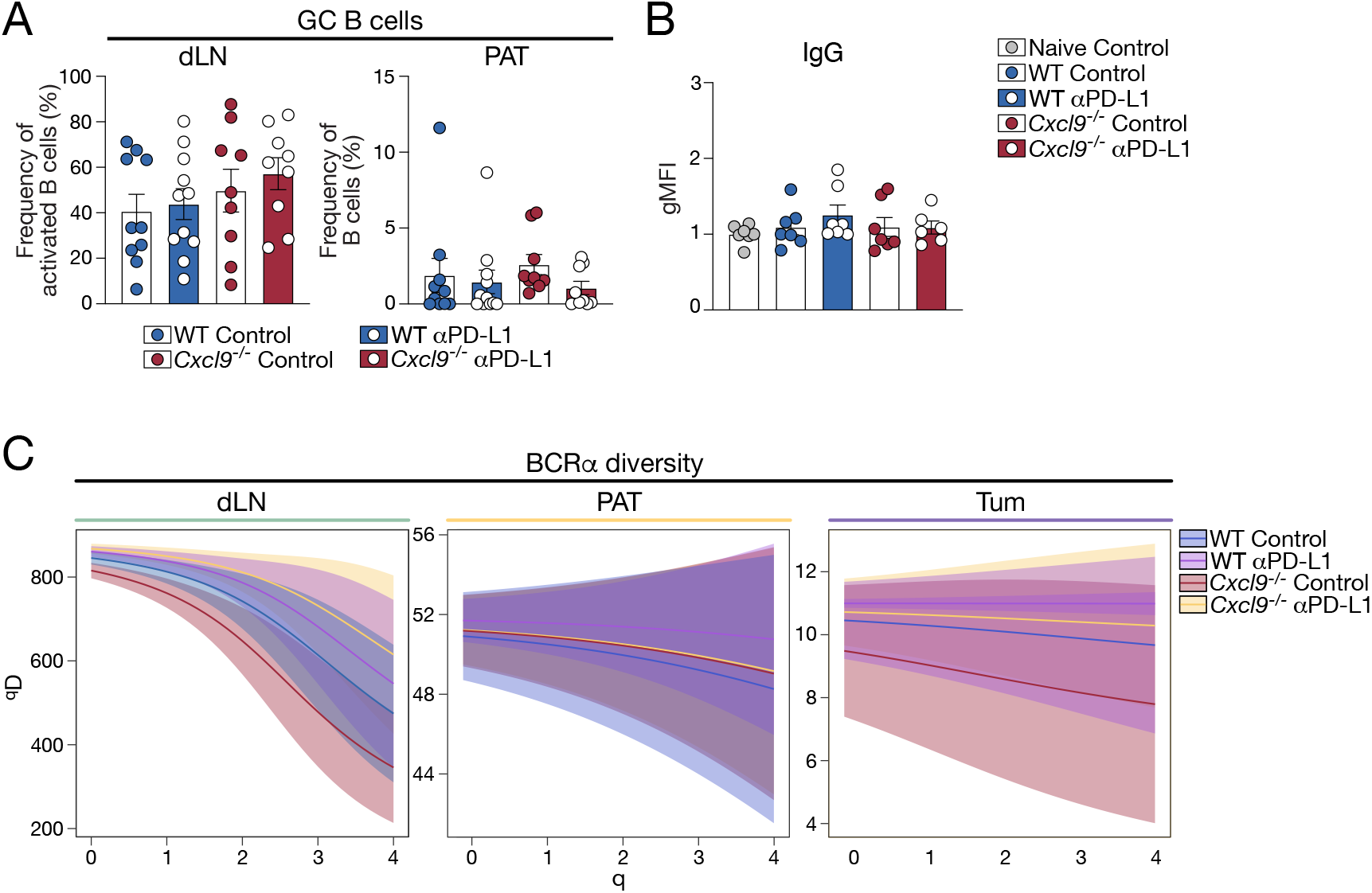
BCR clonal diversity. (A) Frequency of dLN and PAT GC B cell analyzed by flow cytometry. *n* = 9-11 per group. (B) gMFI fold change of serum IgG binding to MC38-OVA^lo^ cells, relative to naïve serum analyzed by flow cytometry. *n* = 6-7 per group. (C) Single cell BCR heavy-chain repertoires analyzed from scRNA-seq. Only annotated B cells carrying a productive IGH contig were included. Data are mean ± SEM. Statistics for (A-B) was determined using two-way ANOVA test.

